# The hybrid nature of task-evoked activity: Inside-out neural dynamics in intracranial EEG and Deep Learning

**DOI:** 10.1101/2020.12.09.417774

**Authors:** Annemarie Wolff, Liang Chen, Shankar Tumati, Mehrshad Golesorkhi, Javier Gomez-Pilar, Jie Hu, Shize Jiang, Ying Mao, Andre Longtin, Georg Northoff

## Abstract

A.

The standard approach in neuroscience research infers from the external stimulus (outside) to the brain (inside) through stimulus-evoked activity. Recently challenged by Buzsáki, he advocates the reverse; an inside-out approach inferring from the brain’s activity to the neural effects of the stimulus. If so, stimulus-evoked activity should be a hybrid of internal and external components. Providing direct evidence for this hybrid nature, we measured human intracranial stereo-electroencephalography (sEEG) to investigate how prestimulus variability, i.e., standard deviation, shapes poststimulus activity through trial-to-trial variability. We first observed greater poststimulus variability quenching in trials exhibiting high prestimulus variability. Next, we found that the relative effect of the stimulus was higher in the later (300-600ms) than the earlier (0-300ms) poststimulus period. These results were extended by our Deep Learning LSTM network models at the single trial level. The accuracy to classify single trials (prestimulus low/high) increased greatly when the models were trained and tested with real trials compared to trials that exclude the effects of the prestimulus-related ongoing dynamics (corrected trials). Lastly, we replicated our findings showing that trials with high prestimulus variability in theta and alpha bands exhibits faster reaction times. Together, our results support the inside-out approach by demonstrating that stimulus-related activity is a hybrid of two factors: 1) the effects of the external stimulus itself, and 2) the effects of the ongoing dynamics spilling over from the prestimulus period, with the second, i.e., the inside, dwarfing the influence of the first, i.e., the outside.

**Significance Statement:** Our findings signify a significant conceptual advance in the relationship between pre- and poststimulus dynamics in humans. These findings are important as they show that we miss an essential component - the impact of the ongoing dynamics - when restricting our analyses to the effects of the external stimulus alone. Consequently, these findings may be crucial to fully understand higher cognitive functions and their impairments, as can be seen in psychiatric illnesses. In addition, our Deep Learning LSTM models show a second conceptual advance: high classification accuracy of a single trial to its prestimulus state. Finally, our replicated results in an independent dataset and task showed that this relationship between pre- and poststimulus dynamics exists across tasks and is behaviorally relevant.

## C. Introduction

The common approach to task-related activity in neuroscientific research has been to investigate the effect of an external stimulus on neural activity and its resulting behavior and cognition. György Buszáki (Buzsáki, 2019) suggests an alternative approach, ‘inside-out’; unlike the traditional ‘outside-in’, it places intrinsic neural activity as a key factor (inside) in modulating activity change related to external stimuli (outside) (see also Northoff et al. 2010, Northoff 2014a and b). Consistent with this approach (Buzsáki, 2019), studies (He, 2013; Baria et al., 2017; Huang et al., 2017; Nieus et al., 2018; Galindo-Leon et al., 2019; Hirschmann et al., 2019; Podvalny et al., 2019) have demonstrated that poststimulus activity levels depend on the initial state, the level of prestimulus activity (Yamagishi et al., 2008; Mathewson et al., 2009; Northoff et al., 2010; Fellinger et al., 2011; Hanslmayr et al., 2013; He, 2013; Milton and Pleydell-Pearce, 2016; Benwell et al., 2017; Huang et al., 2017; Hirschmann et al., 2019). These findings demand the question: how and in what way do prestimulus activity levels shape stimulus-induced activity beyond the external stimulus?

Variability of the signal may be a key factor. As neural responses to the same stimulus show significant variability over trials (Churchland et al., 2010; He, 2013), the quenching of this neural variability occurs after stimulus onset (Churchland et al., 2010; Schurger et al., 2015; Arazi et al., 2017a, 2017b; Haar et al., 2017; Huang et al., 2017, 2018; Daniel et al., 2019). Though much studied, how this neural variability quenching in the poststimulus period - termed trial-to-trial variability (TTV) - is influenced by the prestimulus state is unknown.

TTV describes and indexes (Churchland et al., 2010; He and Zempel, 2013; Ferri et al., 2015; Schurger et al., 2015; Arazi et al., 2017a, 2017b; Huang et al., 2017) the suppression (quenching) of the variability of the spontaneous brain activity by the arrival of the stimulus (Churchland et al., 2010, 2011; He, 2013; He and Zempel, 2013; Dinstein et al., 2015; Arazi et al., 2017a, 2017b; Wolff et al., 2019b). TTV quenching has been observed on multiple levels of neural activity: cellular (Arieli et al., 1996; Monier et al., 2003; Finn et al., 2007; Churchland et al., 2010, 2011; Hussar and Pasternak, 2010; Scaglione et al., 2011; Chang et al., 2012; White et al., 2012; Goris et al., 2014; Mazzucato et al., 2015, 2016; Liu et al., 2016); scalp-level (He and Zempel, 2013; Schurger et al., 2015; Arazi et al., 2017b, 2017a); functional magnetic resonance imaging (fMRI) (He, 2013; Ferri et al., 2015; Huang et al., 2017) (see also (Dinstein et al., 2015) for review of TTV).

In addition to its varying modulation by different stimuli (Churchland et al., 2010, 2011; Hussar and Pasternak, 2010; Arazi et al., 2017b; Wolff et al., 2019b), previous studies suggest that TTV is also dependent on the degree of the brain’s variability at stimulus onset. In non-human data, studies examining prestimulus variability in neural activity have provided direct evidence of its modulating effect on stimulus-related sensory activity (Yamagishi et al., 2008; Romei et al., 2008; Mathewson et al., 2009; Hanslmayr et al., 2013; Luczak et al., 2013; Lin et al., 2015; Scholvinck et al., 2015; Benwell et al., 2017; Gulbinaite et al., 2017; Hennequin et al., 2018; Shimaoka et al., 2019; Huang et al., 2019). Moreover, cell-level studies have shown a strong dependence of poststimulus TTV and behavior (reaction times) on prestimulus variability (Kisley and Gerstein, 1999; Curto et al., 2009; Schurger et al., 2010; Pachitariu et al., 2015). How stimulus related TTV is shaped by prestimulus variability in humans, though, is unknown. We therefore asked what is the electrophysiological relationship in humans between pre- and poststimulus variability as measured with TTV? Closing the loop between these two factors - prestimulus (‘inside’) and external stimulus (‘outside’) - to demonstrate the hybrid nature of stimulus-evoked activity is the aim of the present study using human intracranial stereo-electroencephalography (sEEG).

## Results

### i. Summary of methods and hypotheses

In this study, we aimed to investigate how prestimulus activity shapes poststimulus activity in a hybrid way, being composed of internal and external features whose interaction is supposedly mediated by variability. In addressing this question, we encountered the methodological challenge of linking the continuous ongoing dynamics of the prestimulus period to the measurement of discrete, discontinuous activity time-locked to a stimulus (see (Huk et al., 2018) Huk et al, 2018). To combine both, we therefore tested whether prestimulus temporal SD, as measured in a continuous way through standard deviation, influences poststimulus variability, measured in a discontinuous way by TTV (Huk et al., 2018) (Figure 1).

**Figure 1:**
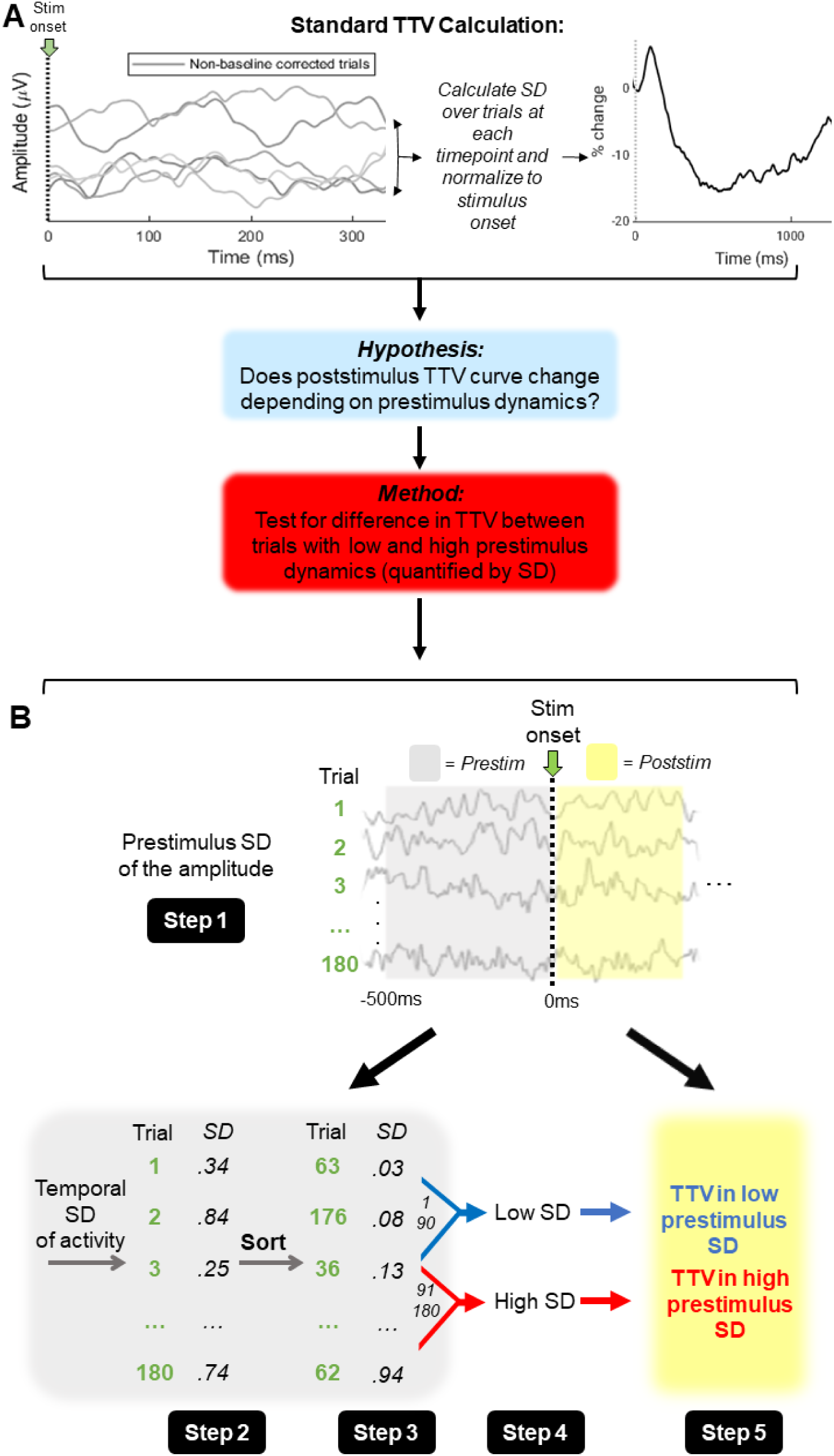
Standard trial-to-trial variability (TTV) calculation and our adjusted method. **A)** TTV is usually calculated by determining the standard deviation (SD) at each timepoint over all trials, then normalized by subtracting and dividing by the SD at stimulus onset (time = 0ms). From this standard calculation, our hypothesis and method of the study resulted. **B)** Calculation of prestimulus SD and sorting of trials into prestimulus low and high. ***Step 1*** – A window prior to stimulus onset (500ms in the broadband) was chosen for each frequency band. ***Step 2*** - The standard deviation of the signal amplitude was calculated in each trial. This continuous measure (Huk et al., 2018) yielded one value per trial. ***Step 3*** – These values were then sorted in ascending order. ***Step 4*** – As there were 180 trials, after sorting trials 1 to 90 were categorized as prestimulus low SD, while trials 91 to 180 were categorized as prestimulus high SD (median split). ***Step 5*** – In each group - prestimulus low, prestimulus high - which consists of 90 trials, TTV was calculated according to the methods of (Wolff et al., 2019b). Hereafter, prestimulus low denotes the TTV computed on the trials with the lower prestimulus SD (trials 1-90) while prestimulus high denotes TTV computed on the trials with the higher prestimulus SD (trials 91-180). TTV is a discontinuous measure (Huk et al., 2018) as it is calculated over trials.

This was done by combining intracranial electroencephalography (sEEG) with Deep Learning neural network models. We investigated intracranial electrophysiological activity, which measures local field potentials (LFP) (Buzsáki et al., 2012), as acquired in a distinct sEEG dataset comprised of 20 human participants. We applied a simple paradigm with two different stimuli and no behavioral response – a no-report paradigm (Tsuchiya et al., 2015). This allowed us to test the impact of the stimulus alone on stimulus-related activity, independent of any behavioral constraints and uncontaminated by any response-related neural activity.

In this sEEG data, we first calculated the standard deviation (SD) (an index of variability) of the signal amplitude in the prestimulus period (varying interval lengths, see methods) (Figure 1). After a median split, trials were assigned to either the prestimulus low or high SD group (median split). TTV was then calculated in each group in the period after stimulus onset, and the area under the curve was measured.

We then sought to examine the effect of the stimulus, including the timing of its effects, on the ongoing pre-poststimulus variability. This was done by comparing the TTV in real trials to that in pseudotrials. Pseudotrials describe time periods between stimulus presentation when a stimulus is absent (Dinstein et al., 2015), and have also been referred to as surrogate trials (He, 2013). Used to model the ongoing dynamics of the spontaneous activity, pseudotrials serve as a baseline for the recorded activity when a stimulus was presented (Huang et al., 2017). When the activity during these pseudotrials was subtracted from the activity of the real trials, the difference shows the stimulus-related activity itself, independent of the impact of the ongoing dynamics (Huang et al., 2017). This allowed us to parcel out and distinguish the effects of the external stimulus itself, and those of the prestimulus dynamics, on stimulus-related activity as measured with TTV.

As TTV is calculated over trials, one curve encompassed data from 90 trials. The need for computing over numerous trials in TTV makes it impossible to account for measuring the impact of the ongoing dynamics on the single trial level. We sought to overcome this by analyzing, and independently validating, the results at the single trial level by applying Deep Learning methods using LSTM neural network models (Alhagry et al., 2017). Specifically, we used our data from the poststimulus activity in individual trials to classify single trials as being either prestimulus low or high (based on a median split) and measuring the accuracy of the classification. This was done first in the trials corrected for prestimulus SD, then in the real trials.

Finally, we replicated our findings in a separate EEG dataset with a report task (Wolff et al., 2019b, 2019a). As the task in this dataset required that participants respond behaviorally, we tested whether the shaping of TTV by prestimulus SD is relevant for behavior.

### ii. Prestimulus SD is the same in real and pseudotrials

After preprocessing of the sEEG data (Daitch and Parvizi, 2018; Helfrich et al., 2018), the event-related potentials (ERP’s) (0.1-70Hz) of the real and pseudotrials were visualized (Figure 2A). This was done to confirm the presence of evoked activity in the real trials and its absence in the pseudotrials. Next, TTV was calculated in the same frequency range in real and pseudotrials to again confirm a response (in real trials) and a lack of response to stimulus (in pseudotrials) (Figure 2B).

**Figure 2:**
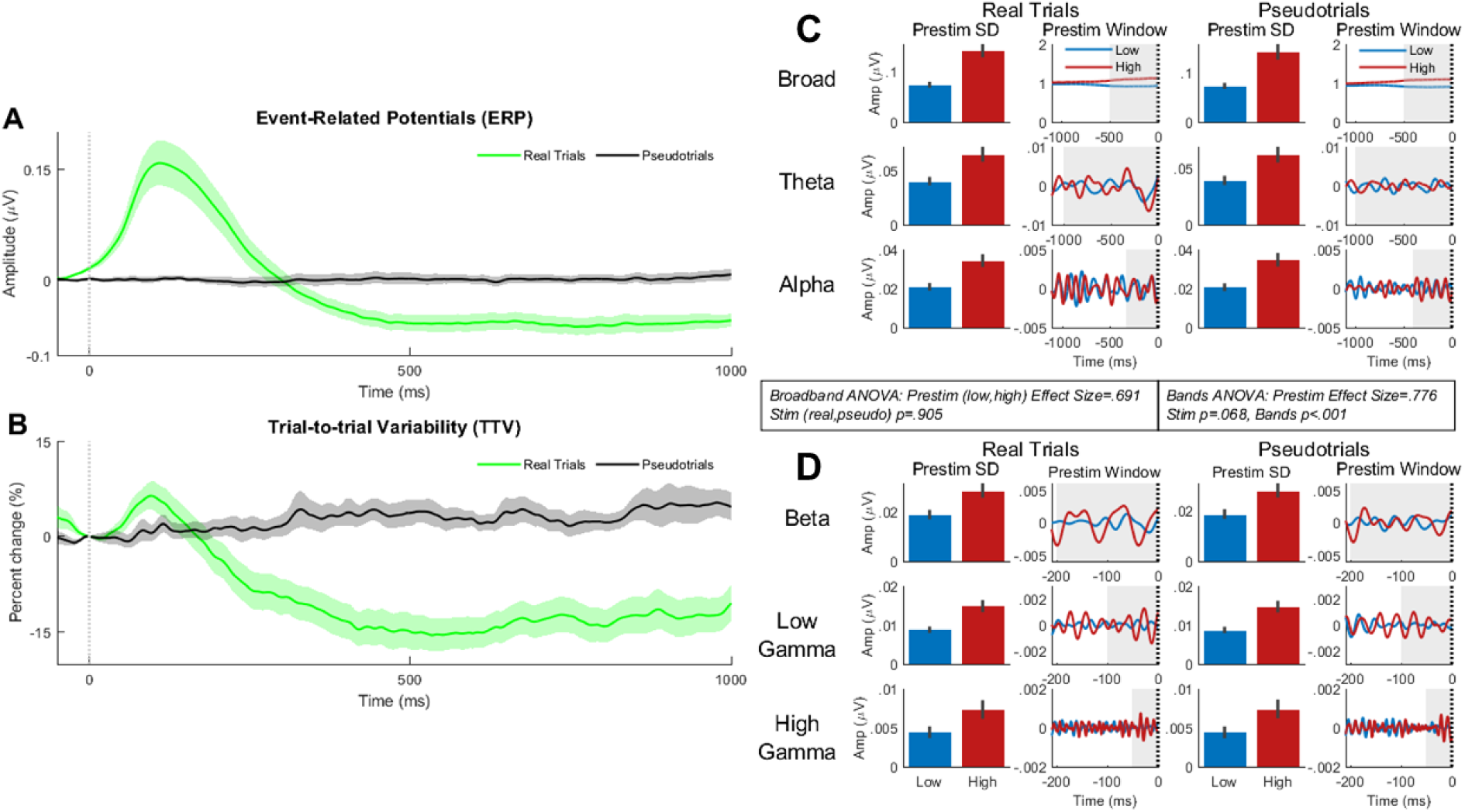
Event-related potentials (ERP), trial-to-trial variability (TTV) and prestimulus SD in real and pseudotrials in all bands. **A)** Event-related potentials (ERPs) for all stimuli in real trials (green) and pseudotrials (black). The ERPs were calculated in the broadband, from 0.1Hz to 70Hz. Shaded areas are standard error. **B)** Trial-to-trial variability (TTV) for all stimuli in real trials (green) and pseudotrials (black) calculated in broadband as the ERPs. Shaded areas are standard error. **C)** Broadband (0.1-70Hz), theta (4-8Hz) and alpha (8-13Hz) prestimulus SD. Prestimulus low (blue) and prestimulus high (red) is shown for real trials (left column) and pseudotrials (right column). The window for calculating the prestimulus SD was 500ms for the broadband, 1000ms for theta, and 400ms for alpha. **D)** For the beta (13-30Hz), low gamma (30-70Hz) and high gamma (70-150Hz) bands, the window of prestimulus SD calculations were 200ms, 100ms, and 50ms respectively. In the broadband, a 2 (prestimulus low, prestimulus high) x 2 (real trials, pseudotrials) repeated measures ANOVA found no effect of stimulus (real, pseudo) (*p*=.905) (prestimulus effect size = .691). In the frequency bands, a 2 (prestimulus low, prestimulus high) x 2 (real trials, pseudotrials) x 5 (bands) repeated measures ANOVA found no effect of stimulus (*p*=.068) and an effect of frequency band (*p*<.001) (prestimulus effect size = .776). Grey shading – time interval of prestimulus SD calculation. Bar and line plots show the mean of all participants. Error bars show standard error.

TTV was defined as the variability changes relative to variability at stimulus onset (see (He and Zempel, 2013; Arazi et al., 2017a, 2017b; Wolff et al., 2019b) for related methods). We calculated the percent change with respect to the value at stimulus onset (He, 2013; Arazi et al., 2017a, 2017b),

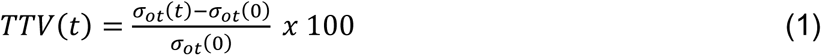

where *σ*_*ot*_(*t*) is the SD of the sEEG signal over trials as function of time *t* and *σ*_*ot*_(0) is the SD over trials at stimulus onset, or 0ms (no difference between SD at stimulus onset between real and pseudotrials with *p*=.065).

As expected, a response to the stimulus was evident in both the ERP and TTV of the real trials (Figure 2A, B). No stimulus-related activity was seen in the pseudotrials. The presence of TTV quenching suggests the potential impact of prestimulus SD and therefore confirmed our use of pseudotrials in the subsequent analyses. TTV quenching thus signified the possible impact of ongoing pre-poststimulus variability on stimulus-related activity in addition to the impact of the external stimulus.

We next grouped the trials according to prestimulus variability. The SD of the prestimulus amplitude was calculated (Figure 1, step 1, 2):

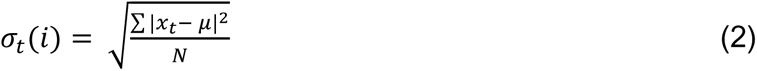

where *σ*_*t*_(*i*) is the SD of the prestimulus interval in trial *i*, *x*_*t*_ is the amplitude at timepoint *t* in the prestimulus interval, *μ* is the mean of the interval and *N* is the number of timepoints in the prestimulus interval. The time interval for this calculation varied according to frequency band (see Methods). Once this was calculated, the SD values were sorted in ascending order and the median value was calculated (Figure 1, step 3). Trials below the median were assigned to the low prestimulus group and those above the median to the high prestimulus group; there were 90 trials in each group.

As the prestimulus median split was based on the SD of signal amplitude in the prestimulus period, we needed to establish if this was the same in real and pseudotrials. In the broadband (0.1-70Hz), a 2 (prestimulus high, prestimulus low) x 2 (real trials, pseudotrials) repeated measures ANOVA found no significant difference between real and pseudotrials (Table 1, Figure 2C).

**Table 1:**
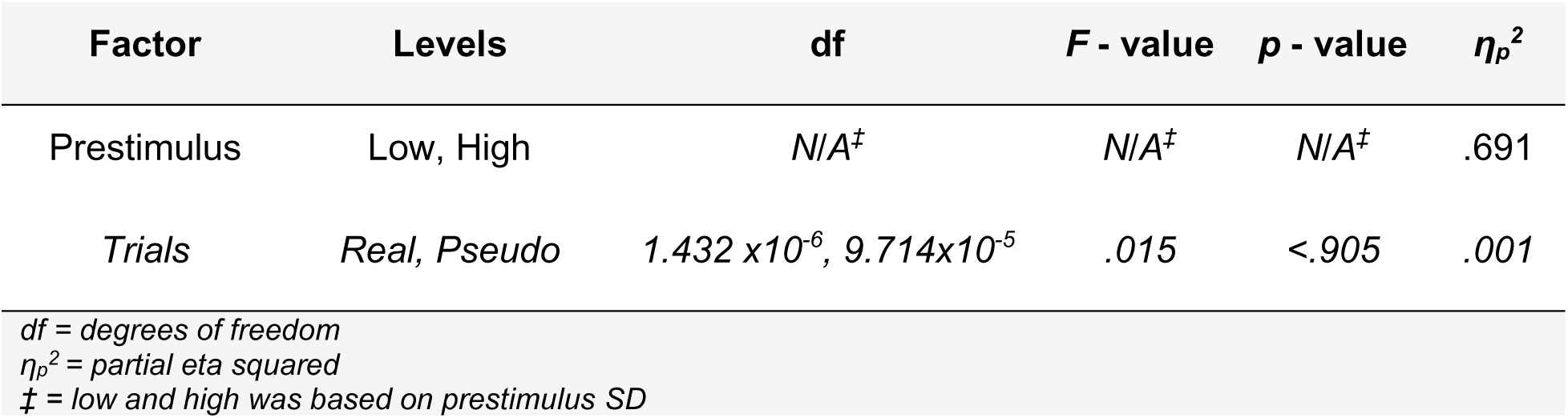
Prestimulus SD 2×2 repeated measures ANOVA results in broadband

| Factor | Levels | df | F - value | p - value | $\eta_p^2$ |
| --- | --- | --- | --- | --- | --- |
| Prestimulus | Low, High | N/A<sup>‡</sup> | N/A<sup>‡</sup> | N/A<sup>‡</sup> | .691 |
| Trials | Real, Pseudo | 1.432 x10<sup>-6</sup>, 9.714x10<sup>-5</sup> | .015 | <.905 | .001 |

df = degrees of freedom
$\eta_p^2$ = partial eta squared
<sup>‡</sup> = low and high was based on prestimulus SD

The repeated measures ANOVA was then done in the different frequency bands, with the bands (theta, alpha, beta, low gamma, high gamma) as an added factor. Again, no significant difference between real and pseudotrials was found (Table 2).

**Table 2:** Prestimulus SD 2×2×5 repeated measures ANOVA results in frequency bands

| Factor | Levels | df | F - value | p – value | $\eta_p^2$ |
| --- | --- | --- | --- | --- | --- |
| Prestimulus | Low, High | N/A<sup>‡</sup> | N/A<sup>‡</sup> | N/A<sup>‡</sup> | .776 |
| Trials | Real, Pseudo | 1,19 | 3.739 | .068 | .164 |
| Bands | theta, alpha,<br>beta, low<br>gamma, high<br>gamma | 1.208,<br>22.961 | 82.858 | <.001* | .813 |

\* = Greenhouse-Geisser corrected
df = degrees of freedom
$\eta_p^2$ = partial eta squared
<sup>‡</sup> = low and high was based on prestimulus SD

To control for the impact of prestimulus mean amplitude, we then did the same median split of the trials and subsequent TTV analysis using the prestimulus mean activity (rather than the prestimulus SD as shown above). We expected that the effect of prestimulus mean (low, high) on poststimulus TTV would no longer be significant. In the TTV AUC for the same 100ms time interval, we found no significant effect of prestimulus mean (*p*=.244) as anticipated. We did find an effect of both stimulus (real trials, pseudotrials) (*p*<.001) and frequency bands (theta, alpha, beta, low gamma, high gamma) (*p*<.001), as we did with the median split according to the SD. Therefore, the difference between real and pseudotrials, and between the frequency bands, was still present when we split the trials according to prestimulus mean, but importantly the difference between prestimulus low and high mean was not significant.

Finally, to control for the different prestimulus time intervals in the different bands in our analysis, we calculated the prestimulus SD again using the same time interval (500ms) for all bands (Supplementary Figures 1 and 2). In both the broadband (prestimulus *p*<.001, stimulus *p*<.001) and the different frequency bands (prestimulus *p*<.001, stimulus *p*<.001, bands *p*<.001), the TTV AUC results were consistent with those found when the prestimulus time interval differed in each band, as we expected. Together, these results show that our poststimulus differences in TTV are related to the variability rather than the mean in the prestimulus interval.

In sum, we established that there was a response to the stimulus in real trials and none in pseudotrials, which verifies our use of the latter. Also, the impact of prestimulus SD showed no significant difference between real and pseudotrials and was higher in the slower bands (than in the faster bands).

### iii. TTV area under the curve showed differences between prestimulus low and high and real and pseudotrials

Once this SD median split had been done, TTV was calculated, as above, over the 90 trials of each group (Figure 1 step 4). To determine if there was a difference in the poststimulus activity in the groups split by prestimulus SD, the area under the curve (AUC) between 450 and 550ms was tested (approximate maximum TTV quenching according to Figure 3A).

**Figure 3:**
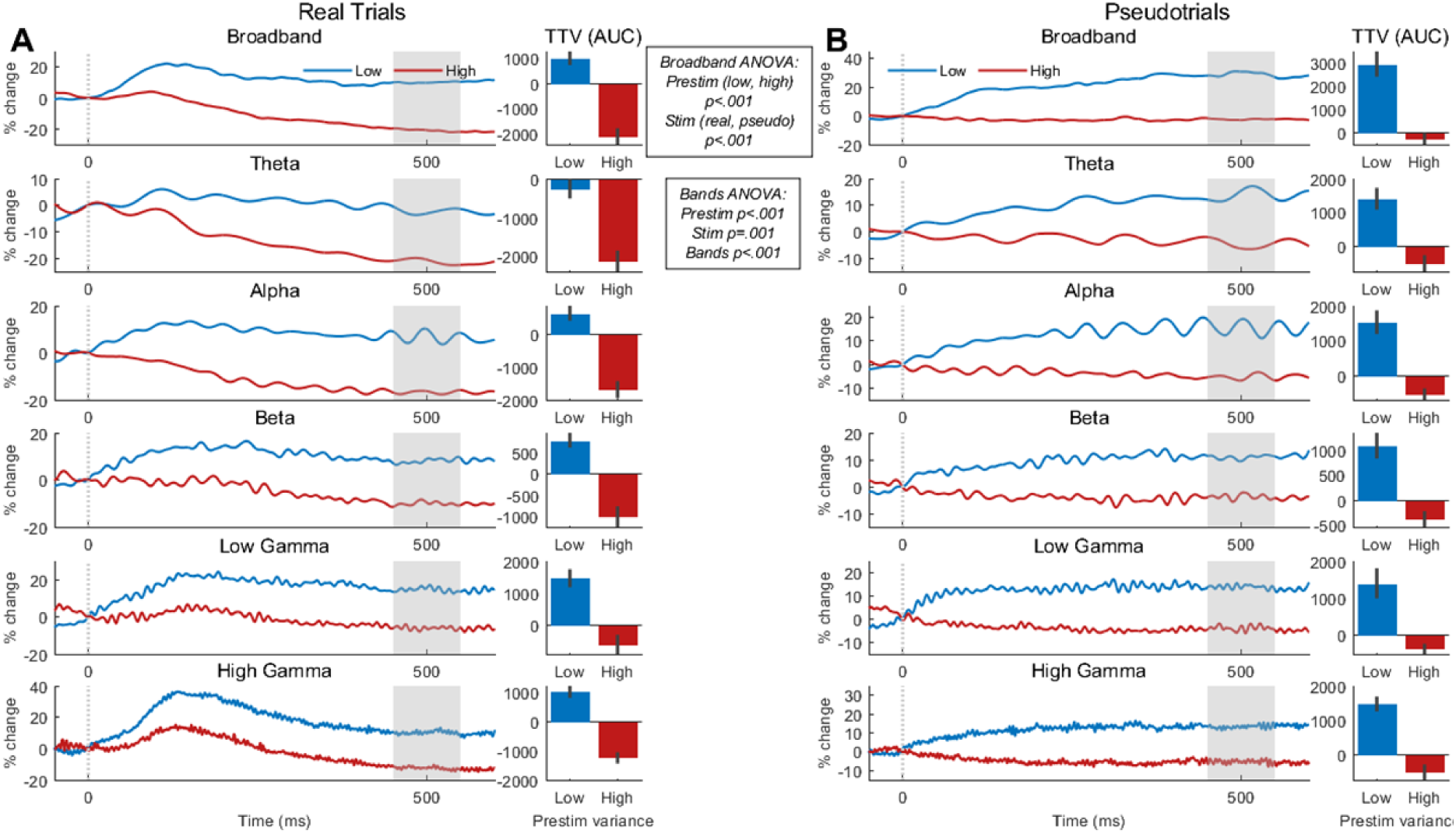
Trial-to-trial variability in real and pseudotrials for all frequency bands. **A)** Trial-to-trial variability (TTV) in real trials for prestimulus low and high. Area under the curve (AUC) from 450 to 550ms was calculated and compared (bar plots). **B)** TTV in pseudotrials with AUC for the same time interval compared. In the broadband, a 2 (prestimulus low, prestimulus high) x 2 (real trials, pseudotrials) repeated measures ANOVA on the AUC found an effect of prestimulus (*p*<.001) and stimulus (*p*<.001). In all bands, a 2 (prestimulus low, prestimulus high) x 2 (real trials, pseudotrials) x 5 (bands) repeated measures ANOVA found effects of prestimulus (p<.001), stimulus (p=.001), and bands (p<.001). Gray shaded areas are interval of calculation of AUC which is shown in the bar plots. Error bars show standard error. Each curve/bar is the mean of all participants.

Again in the broadband, a 2 (prestimulus high, prestimulus low) x 2 (real trials, pseudotrials) repeated measures ANOVA found a significant difference in TTV AUC between low and high prestimulus (*F*(1,19) = 58.692, p < .001 *η_p_^2^* = .755) and between real and pseudotrials (*F*(1,19) = 30.294, p < .001, *η_p_^2^* = .615) (Figure 3). There was a large effect size for both factors.

Next, to measure the same factors in the individual frequency bands, a 2 (prestimulus low, prestimulus high) x 2 (real trials, pseudotrials) x 5 (theta, alpha, beta, low gamma, high gamma) repeated measures ANOVA was done. As in the broadband, there was a significant effect in TTV AUC of prestimulus (*F*(1,19) = 39.288, p < .001, *η_p_^2^* = .674), trials (*F*(1,19) = 14.400, p = .001, *η_p_^2^* = .431) and frequency bands (*F*(2.495, 47.402) = 16.132, Greenhouse-Geisser corrected p < .001, *η_p_^2^* = .459), with a large effect size in all three factors.

After these results, the impact of the interval of the prestimulus SD was examined. To determine if the time interval of measurement for each band had a significant effect on the TTV AUC after stimulus onset, two additional time intervals – one 20% shorter, one 20% longer – were measured (Supplementary Table 1). As the period of each band differs according to their frequency, various prestimulus time intervals were used to measure the prestimulus SD in each frequency band (shorter intervals for higher frequency bands, longer intervals for lower ones). 91% (SD = 1.5%) of the trials in the original and the shorter window were the same, while 91% (SD = 0.8%) of the trials in the original and the longer window were the same. The TTV AUC in the same 100ms interval as above was calculated.

A 2 (prestimulus low, prestimulus high) x 3 (original, shorter, longer window) repeated measures ANOVA found a large effect size of prestimulus in broadband but no significant effect of window on poststim TTV AUC, and a small effect size (Table 3).

**Table 3:** Prestimulus window 2×3 repeated measures ANOVA in broadband

| Factor | Levels | Df | F - value | p – value | $\eta_p^2$ |
| --- | --- | --- | --- | --- | --- |
| Prestimulus | Low, High |  | N/A<sup>‡</sup> | N/A<sup>‡</sup> | .758 |
| Window | Original,<br>Shorter,<br>Longer | 1, 19 | 1.423 | .254 | .070 |

*df = degrees of freedom* *$\eta_p^2$ = partial eta squared* *‡ = low and high was based on prestimulus SD*

Across bands, the same repeated measures ANOVA, with frequency bands as a factor added, again found a large effect size of prestimulus, no effect of prestimulus window, and a significant effect of frequency bands (Table 4). From these results we can conclude that our TTV AUC findings were not determined by our prestimulus interval of measurement.

**Table 4:** Prestimulus Window 2×3×5 repeated measures ANOVA in frequency bands

| Factor | Levels | Df | F - value | p – value | $\eta_p^2$ |
| --- | --- | --- | --- | --- | --- |
| Prestimulus | Low, High | N/A<sup>‡</sup> | N/A<sup>‡</sup> | N/A<sup>‡</sup> | .720 |
| Window | Original,<br>Shorter,<br>Longer | 1.364,<br>25.910 | 3.414 | .064* | .152 |
| Bands | theta, alpha,<br>beta, low<br>gamma, high<br>gamma | 2.705,<br>51.391 | 26.876 | <.001* | .586 |

\* = Greenhouse-Geisser corrected
df = degrees of freedom
$\eta_p^2$ = partial eta squared
<sup>‡</sup> = low and high was based on prestimulus SD

Together our results show that the level of prestimulus SD exerts a strong impact on poststimulus variability in both real and pseudotrials. More generally, our controlled findings show the strong degree to which intrinsic prestimulus SD shapes poststimulus activity, in addition to the effect of the external stimulus.

Since our results above showed a similar difference between prestimulus low and high in pseudotrials as in real trials, we wanted to investigate the effect of the stimulus itself on poststimulus TTV.

Measured neural activity after stimulus onset, A_*m*_, is a sum of multiple activities, plus their interaction (He, 2013; Huang et al., 2017):

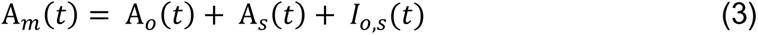

where A_*o*_ is the ongoing spontaneous activity at timepoint *t*, A_*s*_ is the stimulus-related activity, and *I*_*o*,*s*_ is the interaction between the ongoing spontaneous activity and the stimulus-related activity. As it is not possible to measure the interaction between A_*o*_ and A_*s*_ (*I*_*o*,*s*_) directly - A_*o*_ continues to change after stimulus onset (He, 2013) – neural activity was replaced with variability over trials (TTV) in order to isolate stimulus-related activity (A_*s*_). TTV encompasses the interaction of the ongoing spontaneous activity with the stimulus-related activity within it; it is measured relative to SD at stimulus onset and measures the variability over trials. Therefore, to account for this interaction, the neural activity was replaced by the variability over trials, or TTV:

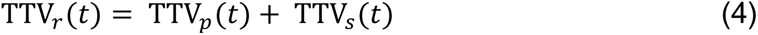

where TTV_*r*_ is the TTV measured in the real trials at timepoint *t*, TTV_*p*_ is the TTV of the ongoing spontaneous activity as measured in the pseudotrials, and TTV_*s*_ is the TTV of the stimulus-related activity (correlations are neglected).

To isolate the effect of the stimulus, the broadband TTV in real and pseudotrials were compared separately for prestimulus low and high (Figure 4A). For each timepoint from stim onset (0ms) to 600ms, two repeated measures *t*-tests were calculated with the respective TTV for all participants. The two tests were a) prestimulus low in real trials compared to prestimulus low in pseudotrials, and b) prestimulus high in real trials compared to prestimulus high in pseudotrials. Therefore, the TTV at timepoint one for all participants (20 patients) in real trials was tested against the TTV at the same timepoint for all participants in pseudotrials. As this was done at each timepoint, it produced a timeseries of *p*-values, as was done previously (He and Zempel, 2013).

**Figure 4:**
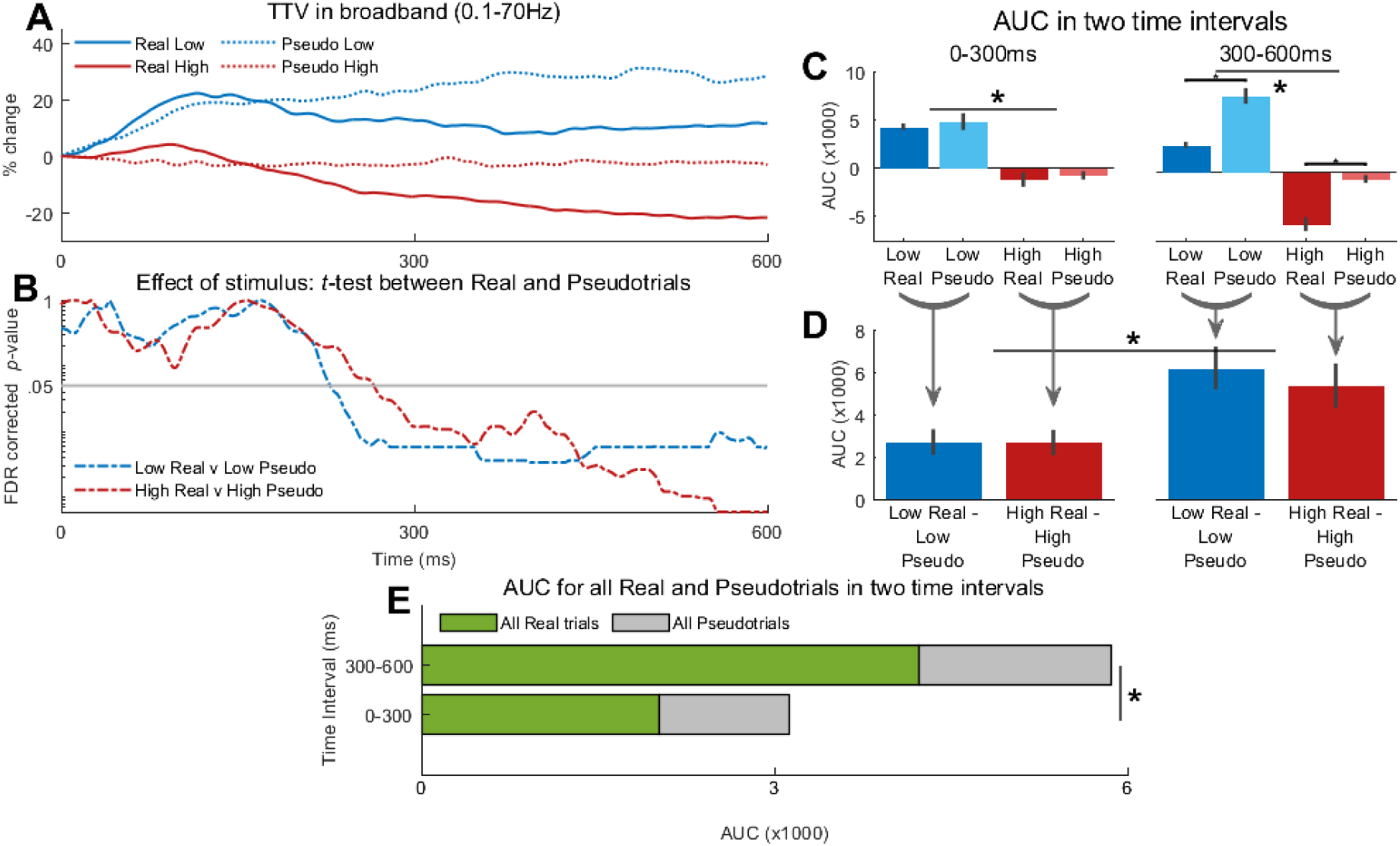
Effect of stimulus on trial-to-trial variability. **A)** Broadband trial-to-trial variability (TTV) in real and pseudotrials. Each curve is the mean of all participants. **B)** To determine the effect of the stimulus, repeated measures *t*-tests were done for all data points between the broadband TTV of real and pseudotrials shown in A. The *p*-values – Benjamini-Hochberg corrected for multiple comparisons – were then plotted for all timepoints. The *p*-values fell below the significance level (.05) just before 300ms. **C)** After the findings in *B*, the poststimulus period was divided into two equal intervals, 0-300ms and 300-600ms. The area under the curve (AUC) was then calculated for all TTV curves in *A* for both intervals. 2×2 repeated measures ANOVAs in each interval found an effect of prestimulus only in the early intervals, and of prestimulus and stimulus in the late interval. **D)** Continuing on from the findings in *C*, for each timepoint the TTV curve for the pseudotrial was subtracted from that of the real trial. The AUC was then calculated and a 2×2 repeated measures ANOVA was done to determine the effect of prestimulus and time interval. No effect of prestimulus was found, though an effect of time interval was. **E)** Finally, in the same intervals from *C* and *D*, the AUC for TTV in all trials – not divided by prestimulus low and high – and pseudotrials was compared. An effect of time interval was found, as was stimulus (real and pseudo). Each bar is the mean of all participants.

This *p*-value timeseries was then corrected for multiple comparisons (Benjamini and Hochberg, 1995) and plotted (Figure 4B). The time interval when the corrected *p*-value timeseries was less than .05 was considered the interval during which the stimulus had an impact. We considered it so as there was a significant difference between the TTV when a stimulus was presented and the TTV when no stimulus was presented; we considered the stimulus to have an impact when there was a difference between the real trials and the pseudotrials. In prestimulus low, this timepoint was found to be at 226ms, while the significance level was passed at 254ms in prestimulus high (Figure 4B).

After visualizing the resulting *p*-value timeseries’ (Figure 4B), the timeseries’ crossed the significance level at slightly before the 300ms mark, or the halfway point of our poststimulus window. As a result, we divided the poststimulus window into two equal intervals (300 timepoints), an earlier one and a later one (henceforth termed ‘early’ and ‘late’).

To determine the effect of prestimulus variability and trials (real, pseudotrials) in these two intervals (early: 0-300ms and late: 300-600ms), the AUC during the two intervals for each of the four TTV curves were compared (Figure 4C). In the early time interval, a 2 (prestimulus low, prestimulus high) x 2 (real trials, pseudotrials) repeated measures ANOVA found a significant effect of prestimulus (*F*(1,19) = 56.291, p < .001, *η_p_^2^* = .748) but not of stimulus (*F*(1,19) = .896, p = .356, *η_p_^2^* = .045). In contrast, the late time interval found a significant effect of both prestimulus (*F*(1,19) = 60.795, p < .001, *η_p_^2^* = .762) and stimulus (*F*(1,19) = 39.402, p < .001, *η_p_^2^* = .675).

Next, to isolate the stimulus-related variability quenching (TTV reduction), equation 4 must be rearranged:

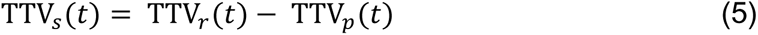

We did this by subtracting the TTV curves at each timepoint *t* in the pseudotrials (TTV_*p*_) from that in the real trials (TTV_*r*_) in the two time intervals (Figure 4D). The AUC of the resulting curves was calculated, and the absolute value was taken (only the magnitude was of interest, not whether the TTV curve increased or decreased in variability). This allowed us to isolate the change in variability due to the stimulus; it is hypothesized that subtracting the pseudotrials effectively removes the variability related to the ongoing activity (He, 2013; Huang et al., 2017). A 2 (prestimulus low, prestimulus high) x 2 (early, late) repeated measures ANOVA found no significant effect of prestimulus SD (*F*(1,19) = .289, p = .597, *η_p_^2^* = .015) and a significant effect of time interval (*F*(1,19) = 19.305, p < .001, *η_p_^2^* = .504).

Finally, in the same early and late time intervals, the TTV AUC for all real and pseudotrials (prestimulus low and high together) was measured (Figure 4E). A 2 (real trials, pseudotrials) x 2 (early, late) repeated measures ANOVA found a significant effect of both stimulus and time interval, and a significant interaction between the two factors (Table 5).

**Table 5:** TTV AUC 2×2 repeated measures ANOVA in broadband for all trials

| Factor | Levels | df | F - value | p - value | $\eta_p^2$ |
| --- | --- | --- | --- | --- | --- |
| Trials | Real, Pseudo |  | 9.649 | .006 | .337 |
| Interval | Early, Late | 1, 19 | 16.601 | .001 | .466 |
| Interaction |  |  | 6.402 | .020 | .252 |

df = degrees of freedom
$\eta_p^2$ = partial eta squared

These findings indicate that the early period of poststimulus activity – 0-300ms - is shaped by both the state-dependent variability of prestimulus SD and the external stimulus. In the later period – 300-600ms - the external stimulus exerts a relatively stronger impact on poststimulus activity than the ongoing spontaneous variability.

### v. Corrected TTV (cTTV) shows greater quenching than TTV (uncorrected)

We next hypothesized that if the TTV curves were corrected for prestimulus SD, the poststimulus differences between prestimulus low and high would decrease, and the magnitude of the TTV quenching would increase. This was tested by calculating TTV corrected using pseudotrials (cTTV),

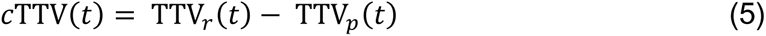

with *TTV_r_* being the curve of the real trials, *TTV_p_* being the curve of the pseudotrials, and *t* being the data point in the timeseries (0 ≤ *t* ≤ 600). We confirmed our hypothesis by measuring the TTV quenching.

The maximum quenching between stimulus onset and 600ms was measured for three groups of trials: 1) all real trials together (180 trials per curve); 2) real trials divided into prestimulus low and prestimulus high (90 trials per curve); 3) corrected TTV (cTTV) – real trials TTV minus pseudotrials TTV - divided by prestimulus low and prestimulus high (90 trials per curve) (Figure 5B). A repeated measures *t*-test (participants provided data to both levels) found a significant difference in the maximum quenching in TTV, but not cTTV (Supplementary Table 2). The difference in quenching between prestimulus low and high disappeared when TTV was corrected for prestimulus SD.

**Figure 5:**
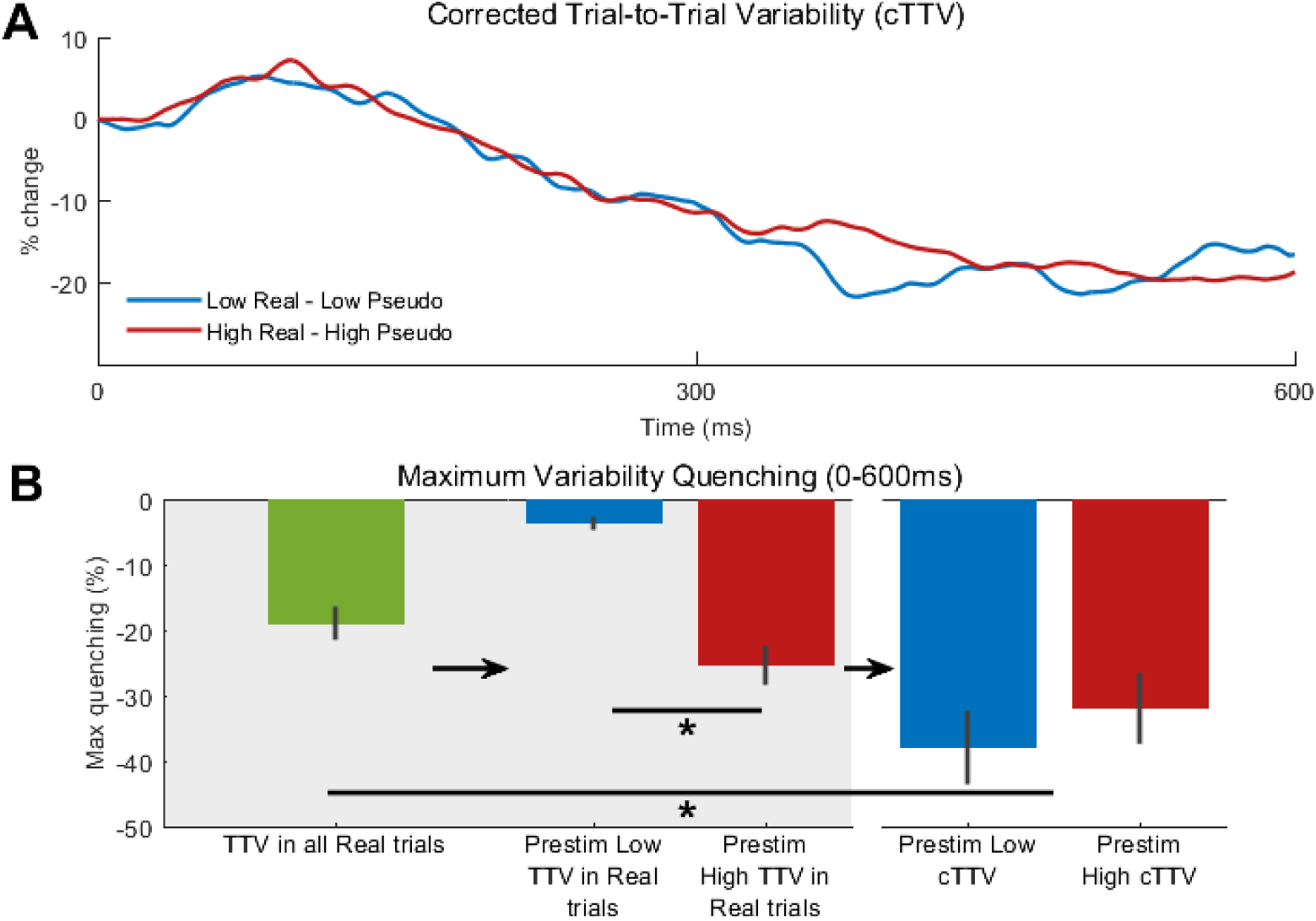
Corrected trial-to-trial variability (cTTV) and its maximum quenching. **A)** Corrected trial-to-trial variability (cTTV) – TTV of real trials minus TTV of pseudotrials – for prestimulus low and high. In these curves the quenching in the latter part of the poststim period reaches approximately 20% for both prestimulus low and high, which contrasts with that of Figure 3A. **B)** TTV maximum quenching (0-600ms) for TTV and cTTV. In all real trials (green bar), max quenching is less than 20%. When the trials are divided into prestimulus low and high (see Figures 1C and 3A), the maximum quenching differs between them. When cTTV is calculated, therefore corrected for prestimulus effects by subtracting pseudotrials, maximum quenching no longer differs between prestimulus low and high, though does differ from that of TTV in all real trials (green bar). Each curve/bar is the mean of all participants.

Lastly, to compare quenching in TTV to cTTV when all trials were combined (180 trials), the maximum quenching in these two curves was measured. A repeated measures *t*-test found a significant difference between TTV and cTTV maximum quenching (Supplementary Table 3), with greater quenching in the cTTV.

In sum, these findings show that correction for prestimulus SD and its ongoing variability yields larger TTV quenching when compared to TTV measured in the standard way, with no correction.

### vi. Single trial analysis - Deep Learning with Long Short-Term Memory (LSTM) neural network showed increased classification accuracy with the real trials

TTV is the SD of activity over trials. Its drawback, therefore, is that it must be calculated over trials and cannot be analyzed at the single-trial level. To overcome this, we used the data from single trials in a Deep Learning neural network. In addition, as our previous analysis had been to look at prestimulus activity’s effect on poststimulus activity, we used the Deep Learning classification to have poststimulus activity predicting prestimulus (low, high) activity.

After considering the results stated above, we asked the following question: Could the trials corrected for prestimulus variability be separated into two groups, prestimulus low and prestimulus high (seen in Figure 5A)? If yes, what would be the accuracy of such a classification? Furthermore, how would the accuracy improve – we hypothesized that it should improve due to the curves we saw in Figure 3A broadband - if the real trials were used to determine the groups (Figure 3A broadband), not the trials corrected for prestimulus variability (Figure 5A)? The maximum quenching values showed no significant difference between these two groups in the corrected trials, but this was not the case in the real trials (Figure 5B).

To measure this, we trained a Deep Learning Long Short-Term Memory (LSTM) recurrent neural network (Alhagry et al., 2017) (Figure 6A) to classify trials as being either prestimulus low or high (Alhagry et al., 2017). This allowed us to independently test our previous analyses, whether ongoing variability affected poststimulus activity.

**Figure 6:**
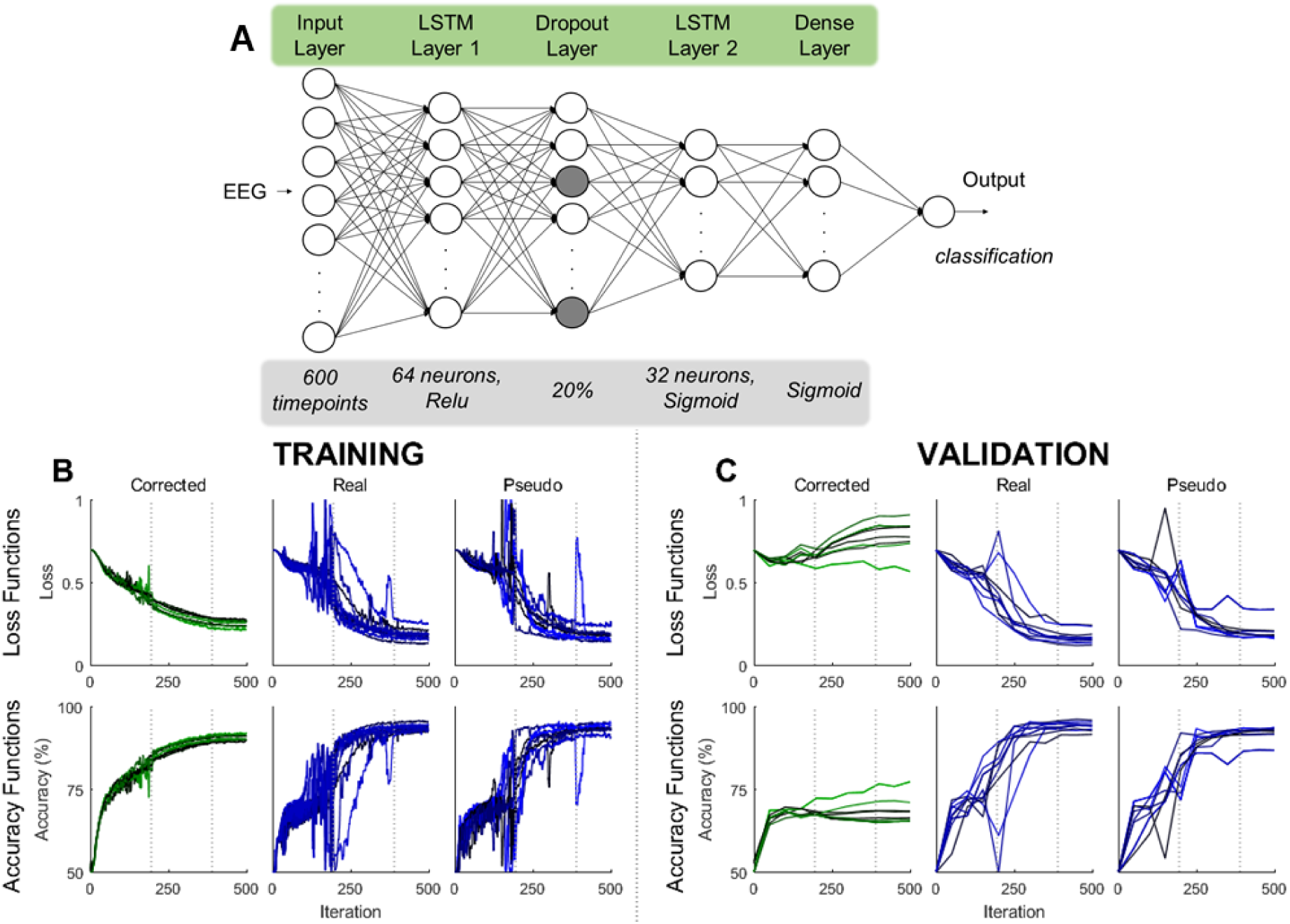
Long short-term memory (LSTM) Deep Learning neural network, training and validation loss and accuracy functions. **A)** LSTM neural network structure according to the methods of Alhagry et al. The dark nodes in the dropout layer signify the randomly assigned 20% dropout per iteration (20% of the nodes were randomly set equal to zero). **B)** Loss and accuracy functions from training of networks for ten repetitions. Vertical dotted lines were the iterations in which the learning rate decreased by 90% as the loss functions increased just before this point in the corrected trials. **C)** Loss and accuracy functions from validation of networks for ten repetitions. Validation occurred every 50 iterations. Vertical dotted lines were the iterations in which the learning rate decreased by 90%.

Also, data at the single trial level were used to increase the number of examples (284 sEEG contacts x 180 trials = 51,120 examples) for the Deep Learning model and to see if the correction at a single trial level yielded similar results to the cTTV results above. Identical to the cTTV correction in equation 5, the corrected trial, *c*T, was calculated as follows:

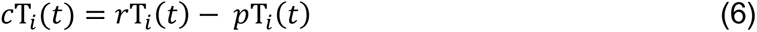

with *r*T being the real trial timeseries, *p*T being the pseudotrial timeseries, *i* denoting the trial number after all trials had been sorted in ascending order according to the calculated prestimulus SD (1 ≤ *i* ≤ 180), and *t* being the timepoint in the timeseries (1 ≤ *t* ≤ 600). Prestimulus variability, on which the trials were assigned to prestimulus low or high groups, was calculated in the broadband from 500ms before the stimulus was presented to stimulus onset (0ms). The timepoints included as inputs to the Deep Learning LSTM models were timepoints 1-600ms. There was, therefore, no overlap in data between the prestimulus calculation and the poststimulus period.

The structure of the LSTM neural network was reproduced from a study which similarly used EEG timeseries data for binary classification (Alhagry et al., 2017). Following the same structure as used there, the timeseries data (sampling rate of 1kHz) were fed into the LSTM neural network via a sequence input layer of 1×600 dimensions (600 data points per trial) (Figure 6A). The sEEG data were not downsampled, nor were it *z*-scored; the preprocessed data used in all previous analyses were input to the model.

From the sequence input layer, the neural network model was arranged as follows (Alhagry et al., 2017): LSTM layer with 64 neurons; Dropout layer with a 0.2 probability (20% of the neurons were randomly set to zero for every iteration as this decreases overfitting); LSTM layer with 32 neurons; Dense layer; classification output layer with a binary cross entropy loss function (measures the performance of the classification, giving a probability of accurate classification) (Figure 6A).

The model was trained on 75% (38,340 examples) of the data while 20% (10,224 examples) was used for validation and the remaining 5% (2,556 examples) was used to test the accuracy. The data split was done randomly using the MATLAB function *dividerand,* and all training, validation and testing was done in MATLAB v2018b using the Deep Learning Toolbox. Finally, ten models with the same structures were trained with each input dataset to ensure consistency of the models and to partially alleviate the modest number of examples. This was measured by the accuracy of classification on the test data.

As shown in Figure 6B and 6C, the first set of Deep Learning models was trained on the corrected trial data (see Figure 6B and 6C for loss and accuracy functions for training and validation for all repetitions). After training ten times, the median binary classification accuracy of the corrected trials was 66% (Mean = 65 ± 2%) (see Supplementary Table 4 and Supplementary Figure 3 for complementary measures) (Figure 6D). Next, to compare this to the uncorrected trials, a model with the same structure was trained ten times on the real trial data – uncorrected for prestimulus variability – and the median accuracy of the classification was found to be 96% (mean = 96 ± 1%). To test for statistical significance, a repeated measures *t*-test found a significant difference between the classification accuracy rates of these two sets of models: *t*(9) = 30.67, *p* < .001.

A final set of models was trained on the pseudotrial data only, again with the identical LSTM structure as done on the previous two sets. After training ten times, the median accuracy of the binary classification was found to be 92% (mean = 93 ± 2%). One final repeated measures *t*-test was done to compare the accuracy of this set to those of the real trials uncorrected for prestimulus variability. This statistical test found a significant difference in the accuracy between these two sets of models: *t*(9) = 3.71, *p* = .005.

Therefore, this classification with LSTM neural networks validated our previous results. It showed much larger differences between poststimulus activity in prestimulus low and high when the data were not corrected for ongoing variability. Furthermore, the accuracy difference between uncorrected real trials and pseudotrials was relatively small but significant. This suggests that the effect of the stimulus is important, though dwarfed by the effect of the ongoing spontaneous activity.

### vii. Replication in EEG: significant effect of prestimulus and trials on TTV area under the curve

To replicate our sEEG findings, analysis was done in a separate EEG dataset with a moral judgement task (see Methods for details) (Wolff et al., 2019b). This task was a report paradigm, in contrast to the no report paradigm of the sEEG, as we needed the response data to examine the behavioral relevance of the prestimulus variability.

After the response of the stimulus on ERP and TTV was determined (Supplementary Figure 4), the area under the curve between 450ms and 550ms was calculated, as was done in the sEEG data (again there was no difference between SD at stimulus onset between real and pseudotrials with *p*=.250). Similarly, in the broadband (0.5-70Hz) a significant effect of both prestimulus (low, high) and trials (real, pseudo) were found in a repeated measures ANOVA (Table 6, Figure 7).

**Figure 7:**
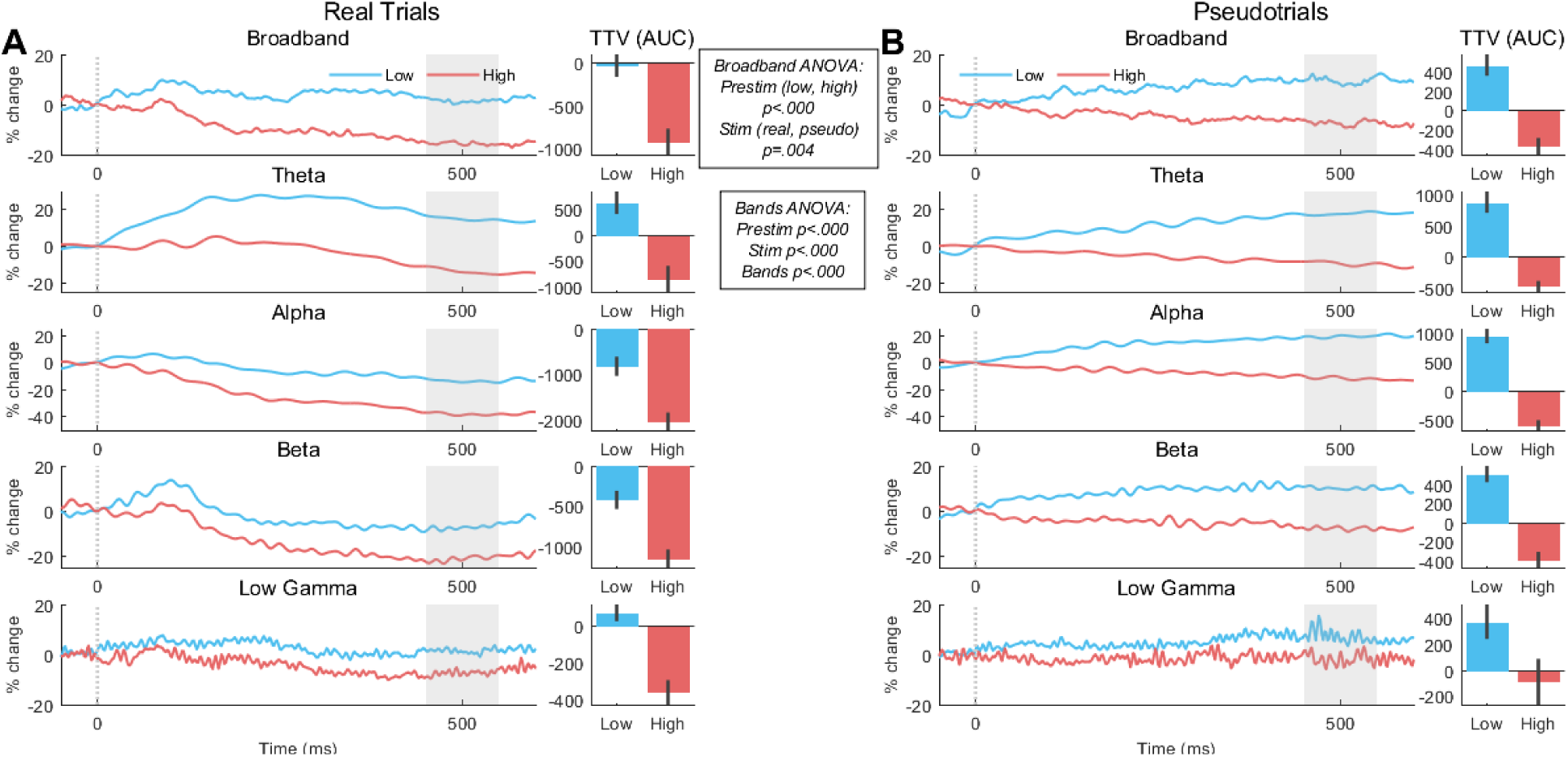
Replication of trial-to-trial variability in real and pseudotrials for all frequency bands in separate EEG dataset. **A)** Trial-to-trial variability (TTV) in real trials for prestimulus low and high. Area under the curve (AUC) from 450 to 550ms was calculated and compared (bar plots). **B)** TTV in pseudotrials with AUC for the same time interval compared. In the broadband, a 2 (prestimulus low, prestimulus high) x 2 (real trials, pseudotrials) repeated measures ANOVA on the AUC found an effect of prestimulus (*p*<.001) and stimulus (*p*=.004). In all bands, a 2 (prestimulus low, prestimulus high) x 2 (real trials, pseudotrials) x 4 (bands) repeated measures ANOVA found effects of prestimulus (p<.001), stimulus (p<.001), and bands (p<.001). Gray shaded areas are the interval of calculation of AUC which is shown in the bar plots. Error bars show standard error. Each curve/bar is the mean of all participants.

**Table 6:** Broadband EEG TTV AUC 2×2 repeated measures ANOVA results

| Factor | Levels | Df | F - value | p - value | $\eta_p^2$ |
| --- | --- | --- | --- | --- | --- |
| Prestimulus | Low, High | 1, 19 | N/A<sup>‡</sup> | N/A<sup>‡</sup> | .785 |
| Trials | Real, Pseudo |  | 10.392 | .004 | .331 |

df = degrees of freedom
$\eta_p^2$ = partial eta squared
<sup>‡</sup> = low and high was based on prestimulus SD

**Table 7:** EEG TTV AUC 2×2×4 repeated measures ANOVA results in frequency bands

| Factor | Levels | Df | F - value | p - value | $\eta_p^2$ |
| --- | --- | --- | --- | --- | --- |
| Prestimulus | Low, High | 1, 19 | 216.39 | <.001 | .912 |
| Trials | Real, Pseudo |  | 27.42 | <.001 | .566 |
| Bands | theta, alpha, beta, low gamma |  | 13.65 | <.001* | .394 |
| * = Greenhouse-Geisser corrected<br>df = degrees of freedom<br>$\eta_p^2$ = partial eta squared | | | | | |

The same was done in the frequency bands, with bands added as a factor (no high gamma in EEG). Again, significant effects of prestimulus (low, high), trials (real, pseudo) and bands (theta, alpha, beta, low gamma) were found in a repeated measures ANOVA (Table 9).

Next, to replicate the cTTV analysis and maximum quenching comparison, the cTTV curves were measured (Supplementary Figure 5A) and the maximum quenching was calculated as above (Supplementary Figure 5B). As in the sEEG results, a repeated measures *t*-test found a significant difference in the maximum quenching in TTV, but not in cTTV (Supplementary Table 5).

Finally, to compare the TTV to cTTV on this measure, the maximum quenching for all trials together was measured. A repeated measures *t*-test found a significant difference on the maximum quenching between TTV and cTTV (Supplementary Table 6).

Together, these findings indicate an analogous relationship of prestimulus SD and poststimulus activity in EEG data recorded on the scalp as in the sEEG data recorded with intracranial depth electrodes. Given that the two datasets had different paradigms, the analogous findings suggest that the shaping of poststimulus activity remains largely independent of cognitive specifics such as stimuli and task. Hence, the observed pre-poststimulus variability shaping may introduce a strong dynamic component into stimulus-related activity.

### viii. Behavioral relevance of prestimulus SD

With the EEG data used for replication, the mean reaction time from the trials with the lowest and highest third (bottom and top 60 trials) of prestimulus variability was calculated for the broadband and each frequency band (Figure 8A).

**Figure 8:**
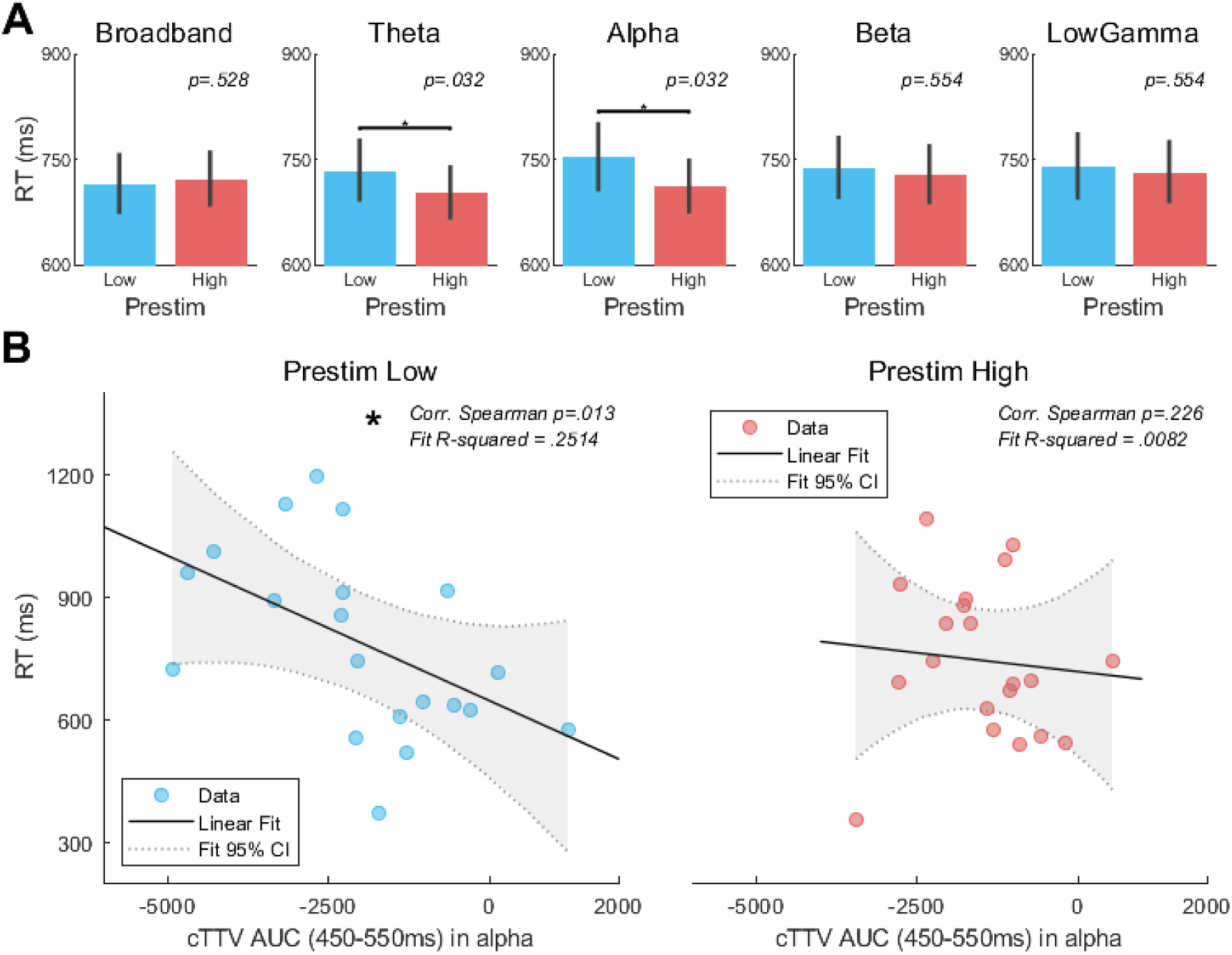
Behavioral relevance of prestimulus SD shown in EEG dataset. **A)** Real trials in each frequency band were split into thirds based on the prestimulus SD and the reaction times for the top and bottom third were extracted. The mean was calculated. A repeated measures *t*-test was done to compare the mean reaction times of prestimulus low and high. There was a significant effect of prestimulus SD in the theta and alpha bands (*p*= .032, .032, Benjamini-Hochberg FDR corrected). Each bar is the mean of all participants. **B)** In these two bands only, the TTV AUC (450-550ms) was Pearson correlated with mean reaction times. In the alpha band, the TTV AUC of prestimulus low had a significant correlation with mean reaction time (*p*=.013), but the prestimulus high did not (*p*=.226).

Repeated measures *t*-tests found a significant difference in the mean reaction times in the theta (*t*(19) = 2.31, *p* = .032) and alpha (*t*(19) = 2.59, *p* = .032) bands. There was no significant difference in the broadband, beta, and low gamma bands (*p* = .528, .554, .494 respectively).

Next, to correlate the standard TTV AUC measured in these two significant frequency bands (theta, alpha) with the mean reaction times, two-tailed Spearman correlations were done. None of the correlations were significant (theta: *p*_low_ = .191 and *p*_high_ = .191; alpha: *p*_low_ = .264 and *p*_high_ = .273).

However, when the same correlations were done between the AUC from the cTTV and the reaction times, the correlation was significant in the alpha band and the prestimulus low group (*r* = -.597, *p*=.013, linear fit R-squared = .2514, linear fit sum of squares due to error = 7.278 x10^5^) (Figure 8B). This was not significant in the prestimulus high group (*r* = -.597, *p*=.013, linear fit R-squared = .0082, linear fit sum of squares due to error = 6.745 x10^5^) or either of the theta correlations (*p*_low_ = .581 and *p*_high_ =.130).

Finally, to test whether 1) the correlation between cTTV AUC and reaction time were significantly different in prestimulus low and high, and 2) whether these correlations were different between cTTV AUC and regular TTV AUC, Fisher’s *r*-to-*z* transformation was done (He et al., 2008). This found that there was a difference between prestimulus low and prestimulus high in the correlation with reaction time and cTTV AUC (*p* = .246), and a difference in the correlations between reaction time of cTTV and TTV (*p*_low_ = .344, *p*_high_ = .936).

In sum, we show that prestimulus SD not only shapes poststimulus activity, but also associated behavior in a complex cognitive task, especially in the alpha band.

## D. Discussion

We investigated the impact of the ongoing dynamics, e.g. prestimulus SD, on poststimulus activity as measured with TTV in both intracranial electrophysiological recordings (sEEG) and scalp-recorded EEG to investigate how prestimulus activity shapes poststimulus activity in a hybrid way. First, we observed that prestimulus SD impacts poststimulus activity in real trials; we observed differences in the latter between prestimulus high and low trials. This served as a basis for our second main finding: the late poststimulus period (300-600ms) showed a greater impact of the external stimulus (relative to the ongoing dynamics) than the early poststimulus period (0-300ms) (where the impact of the ongoing dynamics dominated). Therefore, though both the early and late poststimulus periods where hybrid in their nature, the balance between the internal and external components differed between them. Next, we found that when corrected for prestimulus SD – subtracted the TTV of the pseudotrials from the TTV of the real trials - the maximum quenching of poststimulus variability was the same in trials with low or high prestimulus variability (reflecting the impact of the external stimulus itself), but not when this correction was not done. This indicates the relevance of accounting for prestimulus SD in the analyses of stimulus-related activity when averaging over trials.

To validate and extend our finding of the pre-poststimulus interaction variability on the single trial level, LSTM neural networks were trained – input to models were single trials of poststimulus activity to predict prestimulus group, e.g. high or low prestimulus SD. A significantly higher classification accuracy for prestimulus variability was found when trials of stimulus-related activity were not corrected for prestimulus SD. This shows the importance of accounting for the ongoing dynamics, as indexed by the prestimulus SD, on the single trial level when analyzing stimulus-related activity. We also found that the stimulus itself had a significant effect on the classification accuracy for groups based on prestimulus SD. This shows that, in addition to the ongoing dynamics, the external stimulus itself exerts an important impact, thus demonstrating the hybrid nature of stimulus-related activity. Finally, we replicated all findings in a separate EEG dataset (Wolff et al., 2019b) with a report paradigm (Tsuchiya et al., 2015) that allowed us to show the behavioral relevance of pre-poststimulus variability by shaping reaction time.

Together, the main result is that stimulus-related activity appears to be a hybrid of two components: 1) activity evoked by the stimulus (external source); 2) the ongoing variability carrying over from the prestimulus period to the poststimulus period. This carries major implications for our understanding of stimulus-related activity as we show that the influence of the ongoing variability dwarfs the influence of the stimulus itself.

### i. Poststimulus trial-to-trial variability is dependent on prestimulus variability

Our finding of a difference in TTV between trials with prestimulus low and high variability is consistent with previous studies (He, 2013; Huang et al., 2017) which found that activity at stimulus onset has a differential impact on poststimulus activity. We extend these findings by showing that this difference was found in both the real trials and pseudotrials. The finding is strong evidence of the effect of the ongoing variability on poststimulus activity.

The Deep Learning LSTM neural network models validate – inversely through poststimulus activity predicting prestimulus group - and extends these findings as the models were trained on single trials rather than in our data using TTV (computed over trials). Removal of the prestimulus ongoing variability, as in the corrected trials, showed low accuracy in the classification (66%), though above chance. When the ongoing variability remained – in the real trial poststimulus activity - the classification accuracy increased greatly (96%). The ability of the models to better differentiate between prestimulus low and high when the ongoing variability remains indicates that the prestimulus SD contains a dynamic pattern that shapes stimulus-related activity even on the single trial level; there is a pattern in the data that is easier to detect when trials have not been corrected for prestimulus activity and thus contain this information.

We also observed the impact of the external stimulus itself. The stimulus alone produces a pattern in the data which we also see in the classification accuracy of the corrected trials (above 60%, higher than chance). The important point here, however, is that the impact of the ongoing variability eclipses the impact of the stimulus; the accuracy difference between corrected and real trials is much larger (30%) than the difference between real and pseudotrials (4%). Combined, our empirical findings, and their extension by Deep Learning, show that the ongoing variability, in conjunction with the external stimulus, strongly shape poststimulus activity, even on the single trial level.

### ii. Stimulus has increased effect in later period and prestimulus SD has behavioral relevance

The earlier period (0-300ms) in stimulus-related activity saw a greater influence of the ongoing prestimulus variability than the external stimulus – as seen Figure 4B and 4C. The difference in TTV between low and high prestimulus was significant while the difference between real and pseudo was not. This changed in the later time interval (Figure 4C).

Again, as done above, when the real trials were corrected for prestimulus SD using pseudotrials in this context, the early interval difference between low and high prestimulus disappeared, as it also did in the later interval. This last finding again supports our hypothesis of the substantial impact of the prestimulus SD on poststimulus activity. While the findings in Figure 4E show the different temporal course of both prestimulus SD and external stimulus in shaping stimulus-related activity, the later poststimulus period (300-600ms) shows a larger impact of the stimulus on TTV compared to the earlier period.

Together, we demonstrate that the two components identified as shaping poststimulus-induced activity - ongoing variability and the external stimuli - differ in their temporal course. The ongoing variability exerts stronger effects in the early period while the impact of the external stimulus is stronger in the later time period. It remains to be seen whether the time course of the external stimulus is modulated by diverse stimuli or cognitive requirements related to the said stimulus; this should be a focus of future study.

### iii. Neurophysiological substrate of prestimulus and its ongoing variability

SEEG recordings acquire the activity of local field potentials (LFPs), which measure membrane-potential derived fluctuations in the extracellular space (Buzsáki et al., 2012). Changes in brain LFP’s have been shown to be mediated by synchronization after action potential burst hyperpolarization which can be large and contribute to the extracellular field (Pachitariu et al., 2015).

A relevant study of LFPs in gerbils (Pachitariu et al., 2015) found that in desynchronized cortical states, low frequency fluctuations are suppressed. This allows individual neurons to spike independently with measured activity being more reliable and consistent over trials, and responses to stimuli being faster. In contrast, in a synchronized cortical state the extracellular space shows strong low frequency LFPs fluctuations with high variability in activity over trials and slower responses to stimuli.

These population-level results are consistent with ours on a more macroscale. What they described as the desynchronized state corresponds to our high prestimulus variability; it is associated with stronger poststimulus TTV quenching over trials and faster reaction times in the theta and alpha bands. Similarly, their synchronized state finds its equivalent in our prestimulus low variability which had less TTV quenching and slower reactions times. Due to such correspondence, we infer that, on a cellular level, our high prestimulus variability possibly reflects a desynchronized state that exhibits a higher degree of independent firing of individual neurons, in comparison to the low prestimulus synchronized state. This inference remains tentative, however, and requires a combined investigation of population firing rates and LFPs in humans.

### iv. Limitations

Firstly, though sEEG electrodes are placed in different locations and regions of the brain as they are dependent upon the pathology of the individual patient (epileptic focus), we here did not explicitly analyse the spatial and regional differences. We could observe regional differences in TTV which were largely in accordance with previous data (He 2013, Huang et al. 2017).

Secondly, it should be mentioned that Arazi *et al* (Arazi et al., 2017a) observed the TTV peak to be slightly earlier (200-400ms) than the TTV peak in the current dataset (500-800ms). Though it cannot be verified here, it may be related to differences in stimulus type and length; short visual stimuli for roughly 100ms compared to verbal stimuli lasting approximately 700ms and long (2s) complex visual stimuli with responses.

Thirdly, the prestimulus SD impact on poststimulus activity may have a more cognitive rather than dynamic interpretation. For instance, one may assume that prestimulus SD reflects the prediction of the upcoming stimulus, consistent with predictive coding (Friston, 2010) and recent results on prestimulus alpha (Cao et al., 2017). However, our sEEG no-report paradigm did not include any self-generation of the stimuli and presented stimuli jittered randomly. This makes any prediction (predictive coding) in the prestimulus period unlikely. Therefore, we assume that the prestimulus SD and its impact on poststimulus TTV are more dynamic than cognitive in nature.

## E. Conclusion

To address how prestimulus SD shapes poststimulus activity, as measured with TTV, in a hybrid way, we analyzed an intracranial sEEG data set during a no-report paradigm and combined that with Deep Learning models to extend our data to the single trial level. Our pre-poststimulus variability analyses found three main findings: 1) greater poststimulus variability reduction, e.g. TTV quenching, in trials which had high prestimulus variability; 2) the effect of the stimulus is higher in the later poststimulus period (300-600ms) than the earlier (0-300ms) where the prestimulus dynamics influence dominates; 3) the accuracy of the LSTM Deep Learning models to classify single trials as either prestimulus low or high increased greatly when trained and tested with real trials (that include the prestimulus dynamic effects) compared to trials corrected for prestimulus SD. These results were replicated in a separate EEG dataset and task, which also found that trials with high prestimulus variability in the theta and alpha bands had faster reaction times.

Our findings show that stimulus-related activity is a hybrid, being composed of a) the effects of the external stimulus (outside), and b) the effects of the internal ongoing spontaneous variability (inside), with the second dwarfing the influence of the first. This strongly supports the novel inside-out approach introduced by Buzsáki (Buzsáki, 2019).

## F. Methods

### i. sEEG data and analyses

#### a) Subjects

Twenty patients (25.13 ± 5.57 years; mean ± SD; 8 female) with drug-resistant epilepsy who underwent sEEG exploration in the department of Neurosurgery of Huashan Hospital (Shanghai, China) from September 2016 to May 2017 were included in this study. SEEG is a long-established invasive evaluation method for patients with drug-resistant epilepsies (Parvizi and Kastner, 2018). All patients had comprehensive presurgical evaluations, including a detailed medical history, scalp EEG, magnetic resonance imaging (MRI), and positron emission tomography (PET) scans prior to sEEG exploration. The choice of the anatomical location of electrodes was based on results from the presurgical evaluation and was made by a team of clinicians independent of the present study.

The study was approved by the research ethics committee of Huashan Hospital, at Fudan University, and all aspects of the study were performed according to their relevant guidelines and regulations. Written informed consent was obtained from the patients, or their guardians, for participation in this study.

#### b) Experimental paradigm

The experimental paradigm presented to the patients was comprised of two auditory stimuli: the subjects’ own name and a paired unknown person’s name (in their native language, Chinese) are spoken by the same familiar voice (Qin et al., 2012; Lipsman et al., 2014; Huang et al., 2018). The duration of each name was less than 800ms. The average Root Mean Square (RMS) decibel level was less than -20dB, with the maximum RMS difference between the two names less than 1dB. During this no-report paradigm, participants were presented with 180 trials in total, 90 with their own name and the same number of another name. The intertrial interval (ITI) between trials was 3-3.75s, randomly jittered by 0.25s – this allowed us to minimize anticipation or prediction effects in our analysis of stimulus-related activity as to focus on its dynamic rather than cognitive components.

#### c) sEEG recording

The surgical plans were made by stereotactic neuronavigational software (iPlan Cranial 2.0, Brainlab AG) and the procedures were carried out as previously documented (Gonzalez-Martinez, 2016). Double dosage enhanced T1 images were obtained to identify the blood vessels first then the Leksell stereotactic frame (Elekta) was applied to localize the coordinates of the electrodes. Intracerebral multiple contacts electrodes (HKHS, Beijing, China), with a diameter of 0.8mm and 8–16 contacts, were applied during the surgery which was performed under local anesthesia. The length of each contact was 2mm with 1.5mm between contacts. Post-implantation CT scans were performed the day after implantation surgery to 1) exclude intracranial hematoma, and 2) localize the location of each contact.

The sEEG signals were recorded on a Nihon Kohden 256-channel EEG system (EEG-1200C) with a sampling rate of either 1 or 2kHz, and hardware filtered between 0.01Hz and 600Hz. The signals during recording were referenced to white matter in the brain which was defined by the clinicians.

SEEG is minimally contaminated by artifacts such as swallowing, eye movement, muscle movement, etc., compared to scalp EEG, so it is not prone to artifacts related to gaze position, electrode offset variability, and movement (Parvizi and Kastner, 2018) (see for instance Arazi et al. 2017a and b for excellent control of those factors in EEG). This makes sEEG an ideal tool to record and measure specific variability measures such as TTV.

#### d) Electrodes localization

Electrodes were auto segmented from post-implanted computed tomography (CT) images (Qin et al., 2017). The post-implantation CT was aligned to preoperative MRI images and contacts were calculated and reconstructed. The locations of these contacts were checked manually with neuronavigational software. Finally, the contacts were labeled based on Freesurfer’s parcellation of the MRI.

#### e) sEEG preprocessing

Before any data analysis was performed, the timeseries of each contact was examined by an epileptologist who classified contacts as either seizure or non-seizure. To start, as the data for some of the participants was recorded at a sampling rate of 2kHz while others were recorded at 1kHz, all the former were resampled to 1kHz using MATLAB’s *resample* function which includes an anti-aliasing filter. The events for the task were then imported, superfluous channels (ECG, EMG, etc) were removed, and the contacts previously determined to be seizure contacts by the epileptologist were also removed.

To follow the preprocessing steps of the literature closely, the same methods of two recent sEEG publications (Daitch and Parvizi, 2018; Helfrich et al., 2018), were followed exactly. In MATLAB (The MathWorks, v2012) and according to the methods of Daitch (Daitch and Parvizi, 2018), we calculated the power spectrum for each contact as well as the mean power over all contacts. If the mean power of a given contact was ≥ ±5 SDs of the mean power across all contacts for a given participant, that contact was removed from the data (median = 2 ± 1.2, range = 0-5).

Next, the remaining contacts were two-way zero-phase FIR notch filtered at 50Hz and harmonics (100Hz, 150Hz), and rereferenced to the mean of all remaining contacts. The signal was then bandpass filtered using a two-way, zero-phase FIR non-aliasing filter in the following bands: 0.1-1Hz, 1-4Hz, 4-8Hz, 8-13Hz, 13-30Hz, 30-70Hz, 70-80Hz, 80-90Hz, 90-100Hz, 100-110Hz, 110-120Hz, 120-130Hz, 130-140Hz, 140-150Hz, 150-160Hz, 160-170Hz, and 170-180Hz.

The instantaneous amplitude for each band was then computed by applying the Hilbert transform (MATLAB function *hilbert*) and taking its modulus. Each datapoint for each contact was then normalized by the mean activity of each contact to partially correct for the 1/*f* nature of electrophysiological signals. The signal for each band was then recombined into one signal by taking the mean of all bands per timepoint per contact.

#### f) Stimulus-responsive contact classification

From the preprocessed data, a high-pass two-way zero-phase FIR filter was applied at 70Hz. This left the signal with high frequency band (HFB) gamma between 70-180Hz. The data was then epoched from -2000ms to 2000ms, with a baseline of -200 to 0ms applied to ERP analysis (Helfrich et al., 2018).

We determined which contacts were stimulus responsive according to the methods of Helfrich (Helfrich et al., 2018). Briefly, a contact was excluded from all subsequent analyses when the average HFB response to the stimulus was below a *z*-score of ±1.5 for 10% (50 samples) of consecutive timepoints between stimulus onset (0ms) and 500ms (median = 125 ± 37, range = 45-182 contacts removed per participant).

#### g) Event-related potentials (ERPs)

Event-related potentials (ERPs) were calculated to confirm a response to the stimuli (or its absence in pseudotrials) and for control analysis (see below). As for all other analyses, the data was epoched from -2000ms to 2000ms and was baseline corrected from -200 to 0ms. The mean of all trials (180) over all contacts was calculated for each participant.

#### h) Prestimulus SD

To address the question of the effect of prestimulus variability on poststimulus activity, both continuous and discontinuous measures (Huk et al., 2018) were used. The SD of the prestimulus amplitude – a continuous measure - was first calculated. The SD was calculated according to equation 2, then SD values were sorted in ascending order – the first trial had the lowest prestimulus SD, the last trial had the highest - and the median value was calculated (Figure 1 step 3). Trials below the median were assigned to the low prestimulus group and those above the median to the high prestimulus group, with 90 trials in each.

The time interval for the prestimulus variability calculation varied according to frequency band due to the various period lengths of each band. They were: broadband = -500 to 0ms; theta = -1000 to 0ms; alpha = -400 to 0ms; beta = -200 to 0ms; low gamma = -100 to 0ms; high gamma = -50 to 0ms. To ensure that this window did not affect the results in the TTV AUC, two alternate windows were also calculated and the TTV AUC computed and statistically tested (Table 1).

#### i) TTV in Different Frequency Bands

The frequencies of the sEEG between 0.1 and 70Hz was selected as the broadband for TTV calculation. Moreover, the continuous signal was decomposed into the following frequency bands using a two-way zero-phase FIR filter: theta = 4-8Hz, alpha = 8-13Hz, beta = 13-30Hz, low gamma = 30-70Hz, and high gamma = 70-180Hz. Afterwards, the filtered signals were epoched. The prestimulus SD, including the median split and high/low grouping, and TTV was computed (as explained above) for each band according to the findings of previous studies (Arazi et al., 2017b; Wolff et al., 2019b). Thus, the filtering of the data was done prior to epoching, and the determination of prestimulus high/low SD groups was done separately for each band.

#### j) Real and pseudotrial TTV

The TTV as calculated above includes both a) the effect of the stimulus on the ongoing variability, and b) the ongoing variability that transfers from the prestimulus interval to the poststimulus period (see equation 3 below). To disentangle stimulus-related effects within TTV from those of the ongoing variability, we calculated ‘pseudotrials’ (Huang et al., 2017; Wolff et al., 2019b). Based on fluctuations of the spontaneous activity, the pseudotrials were selected from the ITI preceding each stimulus; specifically, we marked -1500ms (1500ms before onset of the real stimulus) as the virtual stimulus onset and investigated the subsequent 800ms (same duration as the real trials) period as the pseudotrial. In contrast, the real trials were defined as 0 to 800ms of the stimulus-related epoch. Both real and pseudotrial TTV were calculated in the same way as described above (Results, Equation 1). To determine if the prestimulus variability had a significant effect on the poststim TTV, the AUC between 450-550ms was calculated using the MATLAB function *trapz* which employs the trapezoidal method of integration.

#### k) Deep Learning with Long Short-Term Memory (LSTM) neural network

The Deep Learning approach consisted of a long short-term memory (LSTM) recurrent neural network, and the construction, training, validation and testing were done in MATLAB (v2018b) using the Deep Learning Toolbox. A type of recurrent neural network (RNN), the LSTM has been used for inputs of sequences (Alhagry et al., 2017), especially applied to EEG timeseries data (Alexander et al., 2018; Roy et al., 2019), which is why it was chosen here.

From the 1×600-dimension sequence input layer, an LSTM layer with 64 neurons and a Relu activation function followed. Next was a Dropout layer with a 0.2 probability – 20% of the neurons were randomly set to zero as this decrease’s overfitting in the model – followed by a second LSTM layer with 32 neurons and a sigmoid activation function.

Finally, there was a Dense layer with a sigmoid activation function followed by a classification output layer with a binary cross entropy loss function which measures the performance of the classification, giving a probability of accurate classification.

Further parameters of the LSTM network model were 500 iterations, and an initial learning rate of 0.001 with a drop factor of 0.1 every 195 epochs. The learning rate was decreased as the first several attempts to train the model resulted in the divergence of the training and validation losses beginning at epoch 200; the learning rate was therefore decreased to 10% of the previous level (initial learn rate 0.001 multiplied by drop factor 0.1) just before epoch 200, at epoch 195. The frequency of the validation of the network occurred every 50 epochs (10 validation data points).

The model was trained on 75% of the data, validated on 20% and tested on 5%. Despite our ten repetitions of the training, validation and testing of the models, the data split may have influenced the accuracy of the classification. To address this, we trained, validated and tested the model again with the three groups of data, however the split amounts differed. We trained on 75%, validated on 15% and tested on 10% and found consistent results: classification accuracy rate of 65% in the corrected trials, 92% in the real trials and 90% in the pseudotrials.

The model with the specified data input was trained and tested ten times to ensure consistency of the results.

### ii. EEG data and preprocessing for replication

#### a) EEG paradigm

To replicate the findings in the sEEG data, the same analyses were done in a separate dataset with a moral judgement task (Wolff et al., 2019a) (20 participants, 11 females, mean age = 29.9 ± 11.3 years, range of 19-55 years) in scalp EEG. The data was recorded on Neuroscan’s 64 channel Quik-Cap with parameters detailed previously (Wolff et al., 2019a). Written informed consent was obtained from all participants prior to participation (REB# 2009018, University of Ottawa Institute of Mental Health Research).

The task employed here was the externally-guided decision-making control task explained in detail elsewhere (Wolff et al., 2019b, 2019a). Briefly, participants were presented with a black screen divided vertically by a white line. On either side of the center line, two-dimensional stick people were presented, with the numbers on either side varying. Each stimulus contained a total of twelve people, however the ratio between the left and right sides differed and was presented for two seconds. The ITI’s were jittered at 5s, 5.5s and 6s and randomized. The participants’ task was to judge whether there were more people on the left side of the screen than the right side. Each participant was presented with four stimuli repeated 45 times each, therefore 180 trials in total, the same number as in the sEEG task.

Participants responded by a button press (the YES button was counterbalanced across participants) with the mean reaction time being 761 ± 198ms and a range of 391 to 1111ms.

#### b) EEG preprocessing and pseudotrial insertion

The preprocessing for the EEG data was done in MATLAB (The MathWorks, v2018b) with the Optimization, Statistics and Signal Processing Toolboxes using EEGLAB (Delorme and Makeig, 2004) version 14.

To begin, data was resampled to 500Hz using EEGLAB’s *resample* anti-aliasing function. The data was then low- and high-pass filtered using a two-way zero-phase FIR filter at 0.5Hz and 70Hz respectively (High Gamma was excluded). Data were then visually inspected. Channels flat longer than 5s, those that had less than 0.8 correlation with neighboring channels or line noise ±4SD compared to other channels were removed (mode = 2, range 0 to 3). The data was then referenced to the average of all channels, as was done in the sEEG preprocessing. To remove 60Hz electrical noise activity, EEGLAB function CleanLine, which uses a multi-taper sliding window approach to remove electrical line noise, was used.

Data was epoched similarly to the sEEG data (described above) and independent component analysis (ICA) reduced stationary artifacts with the Multiple Artifact Rejection Algorithm (Winkler et al., 2011, 2014).

Identical to methods previously stated (Wolff et al., 2019b), pseudotrials were inserted into the EEG blocks. This was done in the ITI preceding the real stimulus, specifically the onset of the pseudotrial was set 3.5s prior to the onset of the actual stimulus. This allowed for a minimum of 1s between the pseudo and real trial.

#### c) Reaction time data and behavioral relevance

To determine if there was a significant difference in reaction time between trials with prestimulus high or low variability, the prestimulus SD of trials were split into three groups (triple-split). The corresponding reaction times from the top and bottom third (ascending SD values 1-60 and 121-180) for each frequency band were then extracted and the mean of each group was computed. A repeated measures *t*-test was done for each frequency band to compare the mean reaction times between the low and high prestim.

To correlate the TTV AUC in the difference curves described above (TTV real trials minus TTV pseudotrials), the reaction times for the median split based on the prestimulus SD were extracted and the mean was computed. A two-tailed Spearman’s correlation (some of the distributions were non-normal) between these mean reaction times and the TTV AUC was done for both the prestimulus high and low. To support this correlation, a linear polynomial curve was fit to the data using MATLAB v2018b’s *fit* function (fitType = ‘poly1’). Finally, to contrast these results with those from the TTV curves that were not corrected for prestimulus variability, the same analysis was done with the TTV AUC from the real trials.

### iii. Correlation and Statistics

All statistics were computed either in SPSS v24 or MATLAB v2018b using the Statistics Toolbox, and the significance level for all tests was .05. To control for multiple comparisons, the False Discovery Rate by Benjamini-Hochberg (Benjamini and Hochberg, 1995) was applied to the *p*-values of all statistical tests (See the result sections for more detailed statistics).

Repeated measures *t*-tests and ANOVA’s were used as all participants contributed to both/all levels (real trials/corrected trials/pseudotrials, prestimulus low/prestimulus high, etc). All assumptions of ANOVA’s were met prior to calculating the statistical test.

## G. Acknowledgments

This work was supported by the EJLB-Michael Smith Foundation, the Canadian Institutes of Health Research, the Ministry of Science and Technology of China, the National Key R&D Program of China (2016YFC1306700), the Hope of Depression Foundation (HDRF), and the Start-Up Research Grant in Hangzhou Normal University (to Georg Northoff). This research has also received funding from the European Union’s Horizon 2020 Framework Program for Research and Innovation under the Specific Grant Agreement No. 785907 (Human Brain Project SGA2).

The first author (A Wolff) would like to thank the Sachs Lab - Chadwick Boulay in particular - and the University of Ottawa Brain and Mind Institute for their applied Deep Learning in intracranial neurophysiology workshop; it provided the basis for the Deep Learning aspect of this study.

## H. Author Contributions

LC, SJ, JH, and YM acquired the sEEG data, while YM was responsible for overseeing the patient recruitment and data acquisition. AW, ST and GN conceived of the idea of the study, AW and ST did the preprocessing and analysis of the sEEG data, and AL provided feedback and suggested further analyses. AW and MG performed the LSTM Deep Learning analyses; MG interpreted the Deep Learning results; JGP made suggestions for the Deep Learning section of the manuscript and its figures. AW acquired and analyzed the EEG replication data. AW and GN wrote the manuscript, while ST, JGP and AL edited the manuscript and made suggestions.

## Supplementary Figures

**Supplementary Figure 1:**
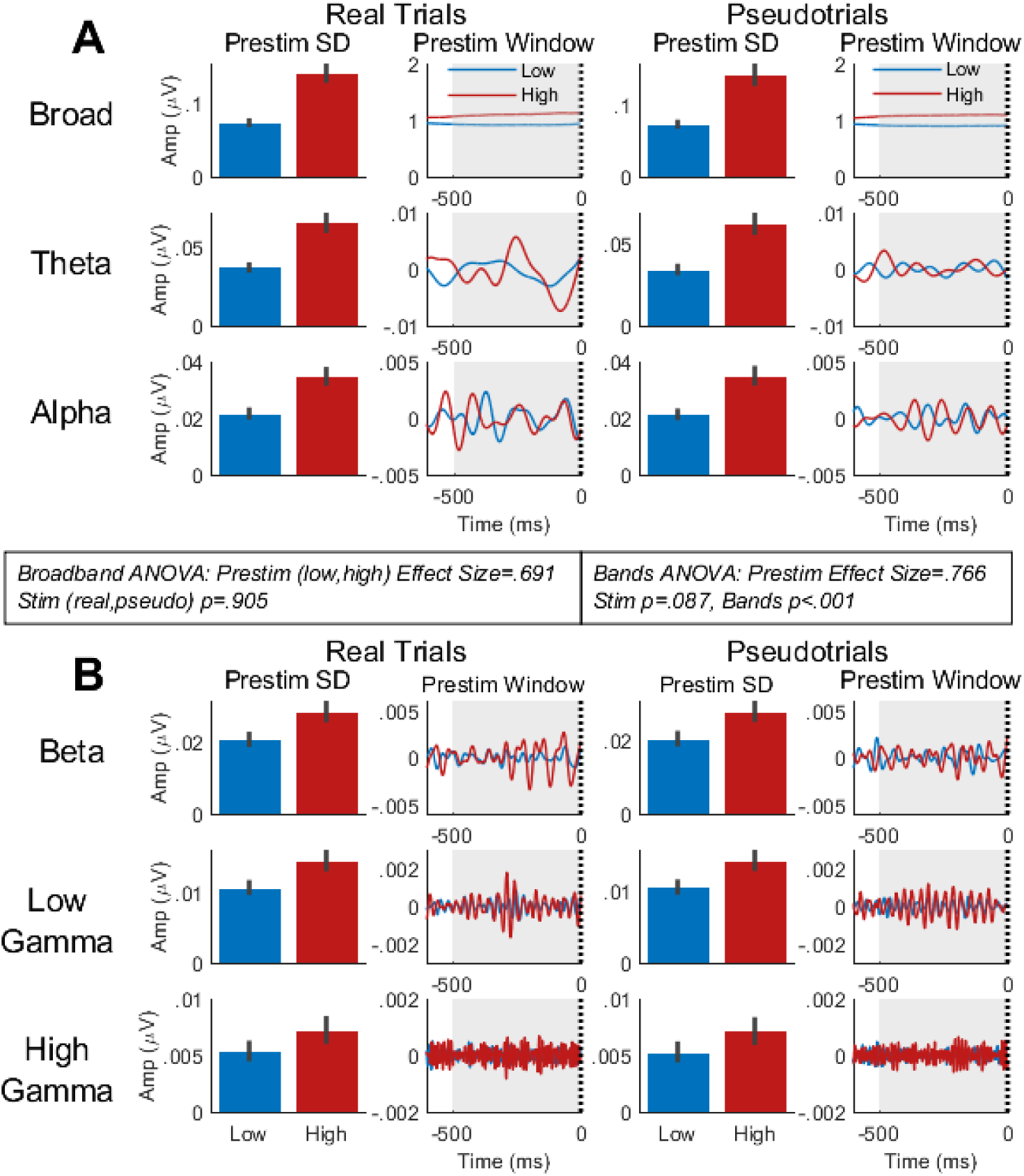
Prestimulus SD calculated during the same 500ms time window in real and pseudotrials in all bands. **A)** Broadband (0.1-70Hz), theta (4-8Hz) and alpha (8-13Hz) prestimulus SD. Prestimulus low (blue) and prestimulus high (red) SD is shown for real trials (left column) and pseudotrials (right column). The window for calculating the prestimulus SD was 500ms for all bands. **D)** For the beta (13-30Hz), low gamma (30-70Hz) and high gamma (70-150Hz) bands, the window of prestimulus SD calculations were 500ms similarly. In the broadband, a 2 (prestimulus low, prestimulus high) x 2 (real trials, pseudotrials) repeated measures ANOVA found an effect size for prestimulus of .691 and no effect of stimulus (real, pseudo) (*p*=.905). In the frequency bands, a 2 (prestimulus low, prestimulus high) x 2 (real trials, pseudotrials) x 5 (bands) repeated measures ANOVA found an effect size for prestimulus of .766, no effect of stimulus (*p*=.087), and an effect of frequency band (*p*<.001). Grey shading – time interval of prestimulus SD calculation. Bar and line plots show the mean of all (20) participants. Error bars show standard error.

**Supplementary Figure 2:**
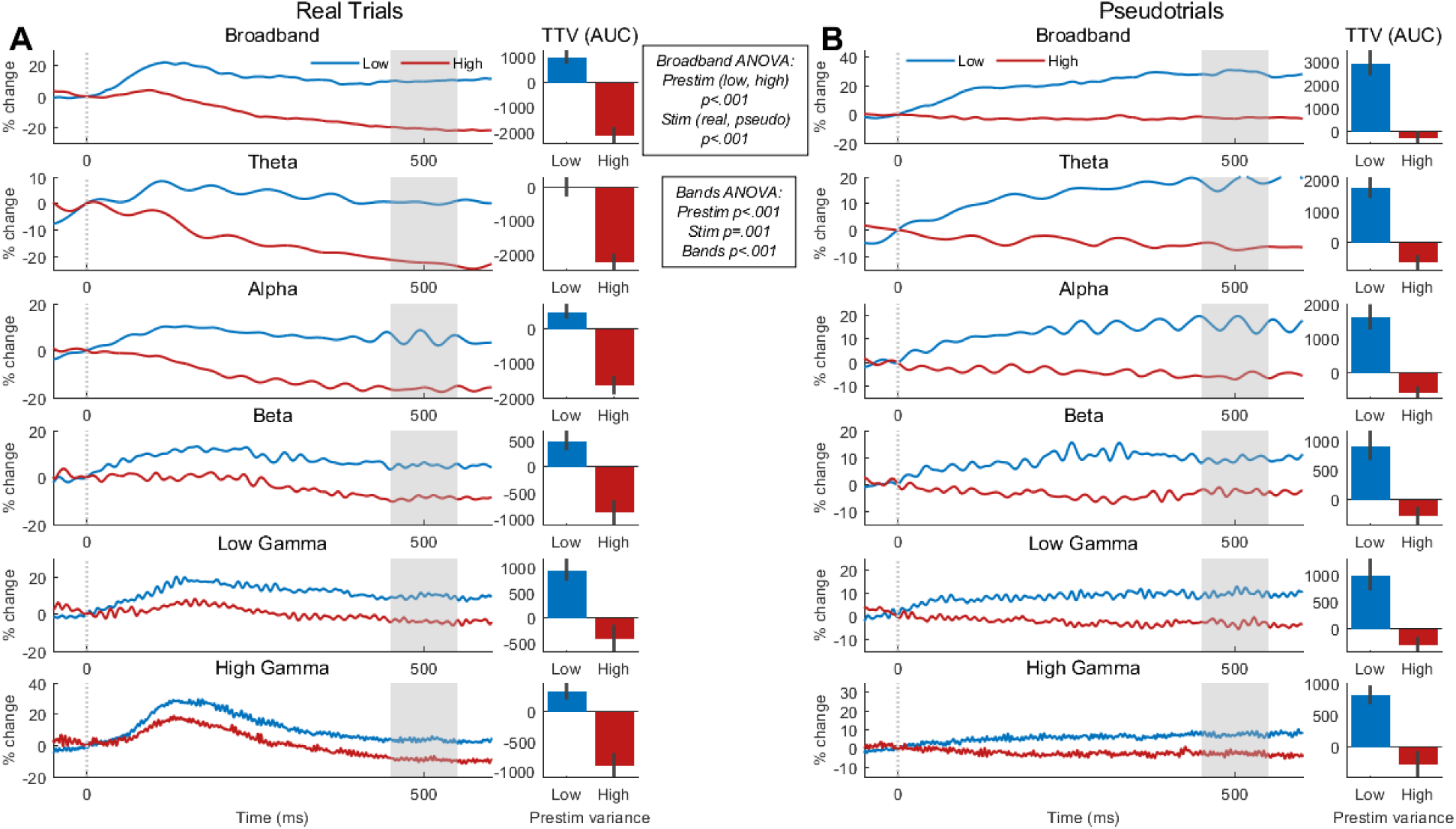
Trial-to-trial variability in real and pseudotrials for all frequency bands when the prestimulus SD was measured in a 500ms window for all bands. **A)** Trial-to-trial variability (TTV) in real trials for prestimulus low and high. Area under the curve (AUC) from 450 to 550ms was calculated and compared (bar plots). **B)** TTV in pseudotrials with AUC for the same time interval compared. In the broadband, a 2 (prestimulus low, prestimulus high) x 2 (real trials, pseudotrials) repeated measures ANOVA on the AUC found an effect of prestimulus (*p*<.001) and stimulus (*p*<.001). In all bands, a 2 (prestimulus low, prestimulus high) x 2 (real trials, pseudotrials) x 5 (bands) repeated measures ANOVA found effects of prestimulus (*p*<.001), stimulus (*p*=.001), and bands (*p*<.001). Gray shaded areas are interval of calculation of AUC which is shown in the bar plots. Error bars show standard error. The curves shown are the mean of all (20) participants.

**Supplementary Figure 3:**
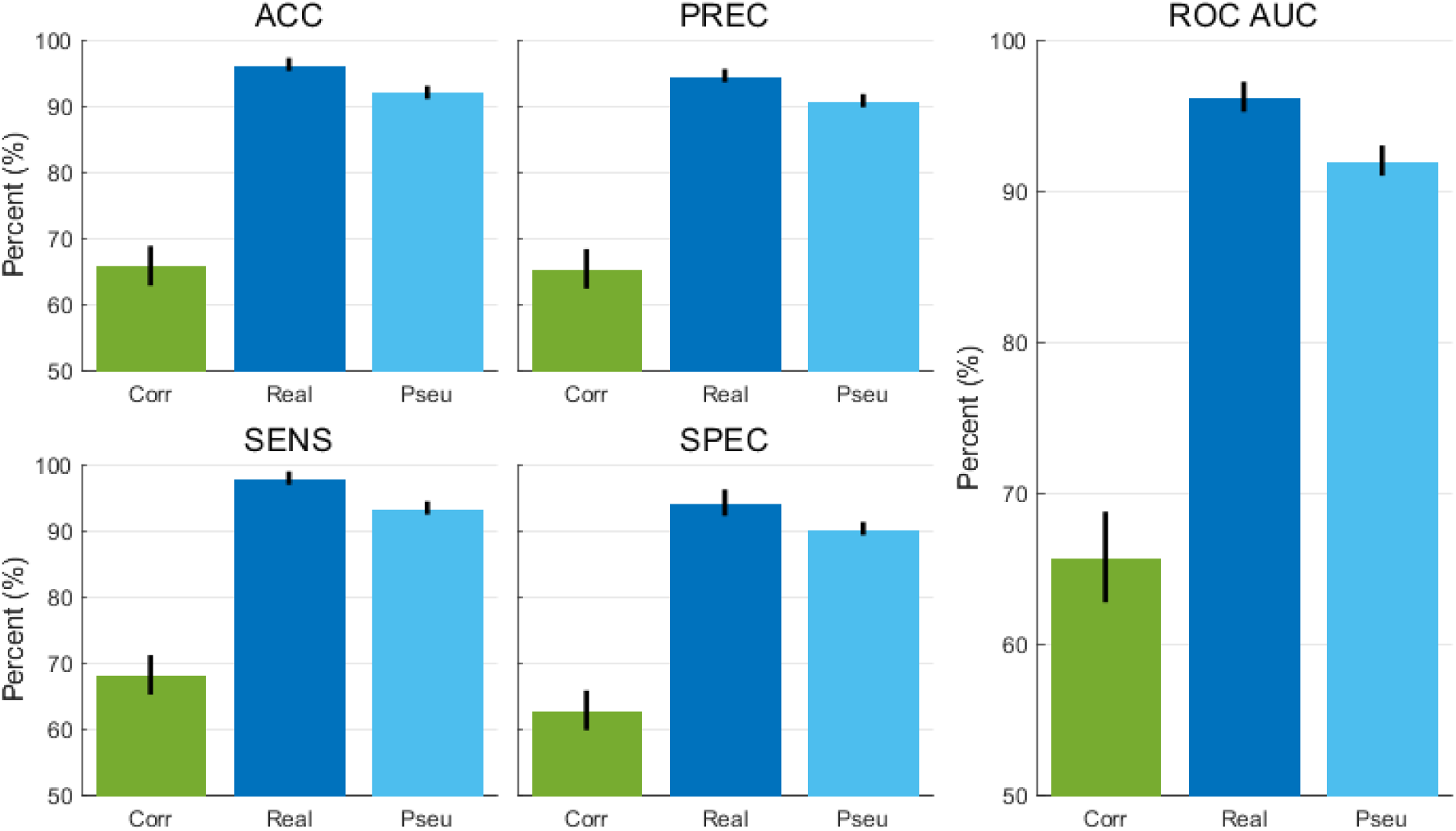
Long short-term memory (LSTM) Deep Learning neural network testing results. The bars represent the median percentages for the three groups of models for each measure, with the error bar denoting the standard deviation over the ten repetitions of the training and testing. For each measure (for equations and values see Table 9), the median of the corrected trials was the lowest, the real trials the highest, and the pseudotrials between them but closer to the real trials. ACC = Accuracy; PREC = Precision; SENS = Sensitivity; SPEC = Specificity; ROC AUC = Receiver operating characteristic curve area under the curve.

**Supplementary Figure 4:**
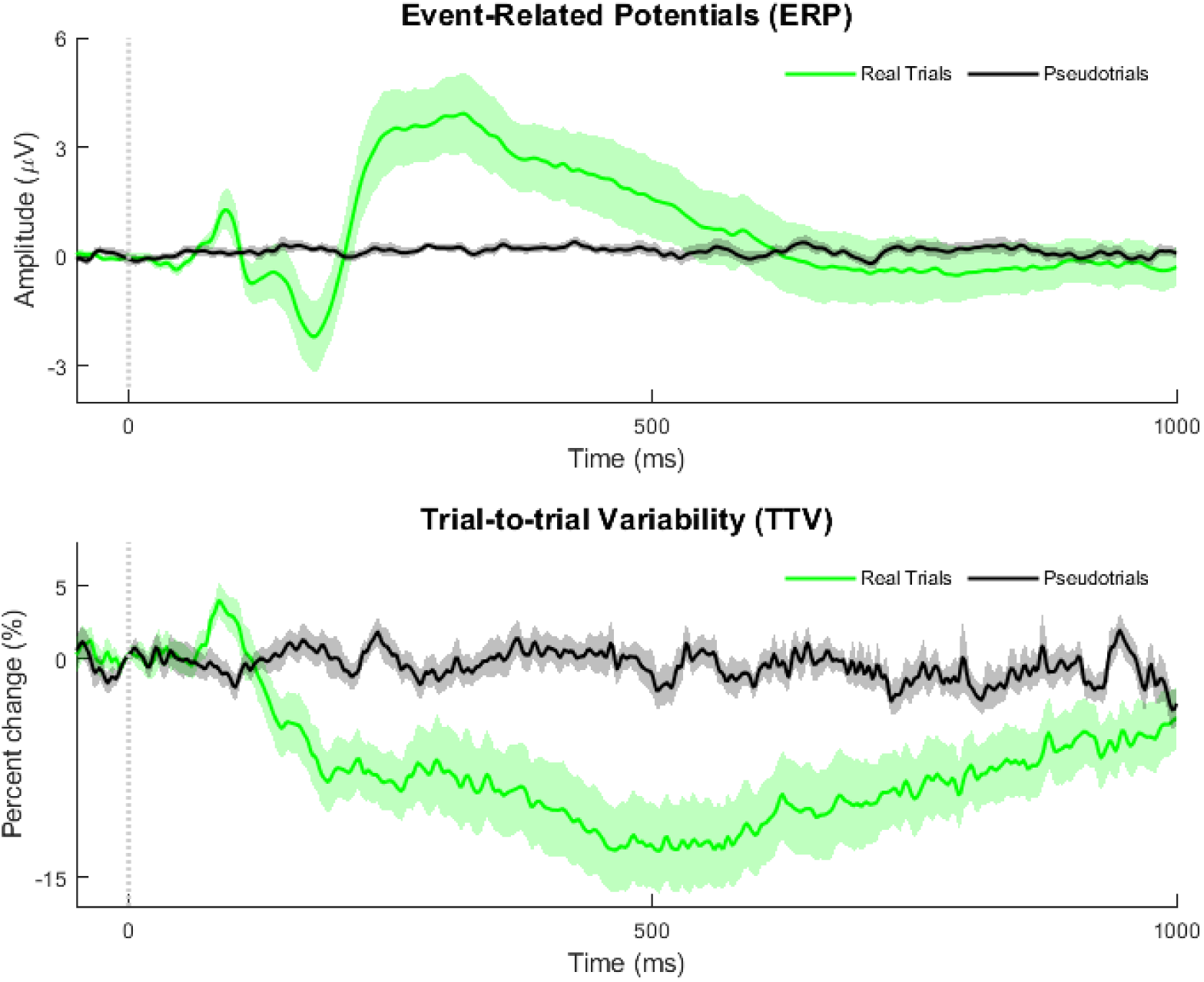
Replication of event-related potentials (ERP) and trial-to-trial variability (TTV) in separate EEG dataset. Top row: Event-related potentials (ERPs) for all stimuli in real trials (green) and pseudotrials (black). The ERPs were calculated in the broadband, from 0.5Hz to 70Hz. Shaded areas are standard error. Bottom row: TTV for all stimuli in real trials (green) and pseudotrials (black) calculated in broadband as the ERPs. Shaded areas are standard error.

**Supplementary Figure 5:**
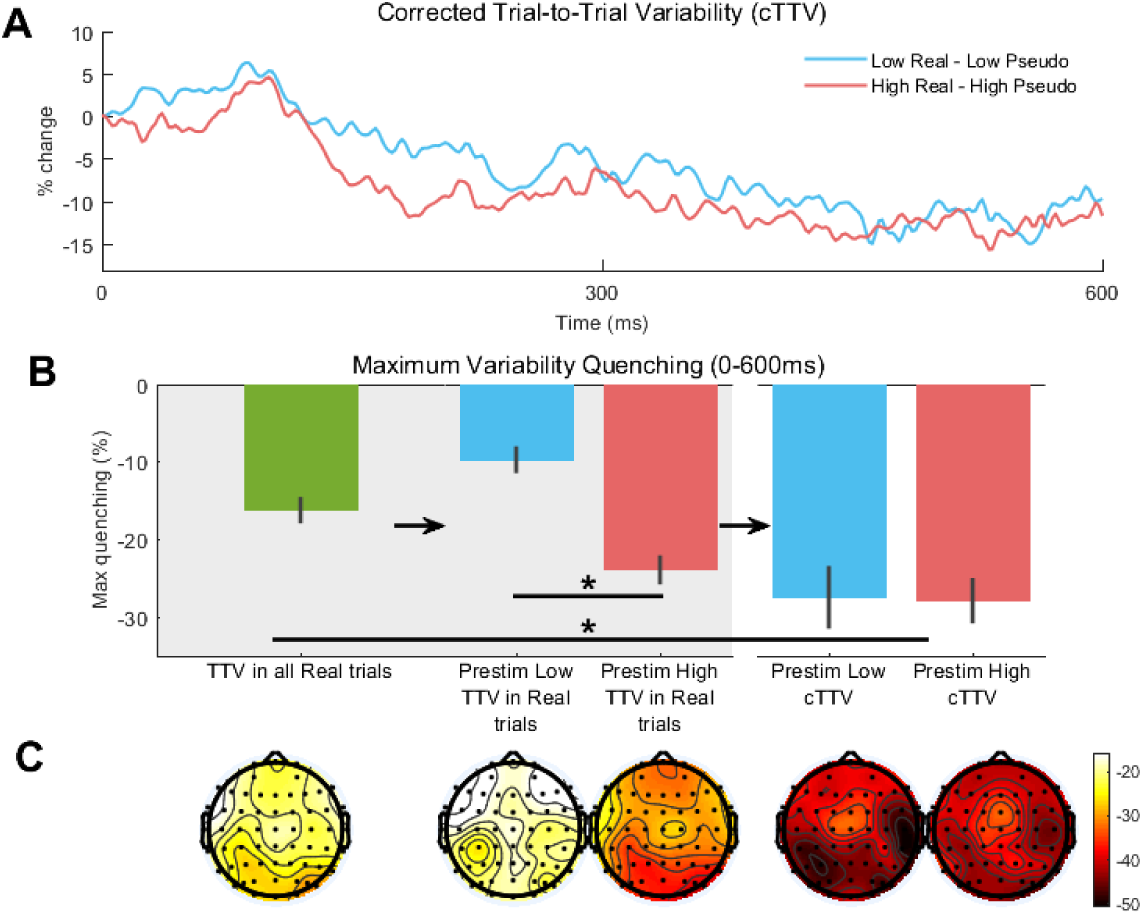
Replication of corrected trial-to-trial variability (cTTV) and its maximum quenching in separate EEG dataset. **A)** Corrected trial-to-trial variability (cTTV) – TTV of real trials minus TTV of pseudotrials – for prestimulus low and high. In these curves the quenching in the latter part of the poststim period reaches approximately 15% for both prestimulus low and high, which contrasts with that of Figure 6A. **B)** TTV maximum quenching (0-600ms) for TTV and cTTV. In all real trials (green bar), max quenching is less than 20%. When the trials are divided into prestimulus low and high (see Figures 1C and 6A), the maximum quenching differs between them. When cTTV is calculated, therefore corrected for prestimulus effects by subtracting pseudotrials, maximum quenching no longer differs between prestimulus low and high, though does differ from that of TTV in all real trials (green bar). **C)** Topographical maps of maximum quenching during the time interval of 0-600ms.

**Supplementary Table 1:** Prestimulus windows of measurement for each frequency band

| Frequency Band | Prestimulus window start prior to stim onset |  |  |
| --- | --- | --- | --- |
|  | Original (ms) | Shorter (ms) | Longer (ms) |
| broadband | 500 | 400 | 600 |
| Theta | 1000 | 800 | 1200 |
| Alpha | 400 | 320 | 480 |
| Beta | 200 | 160 | 240 |
| low gamma | 100 | 80 | 120 |
| high gamma | 50 | 40 | 60 |

**Supplementary Table 2:**
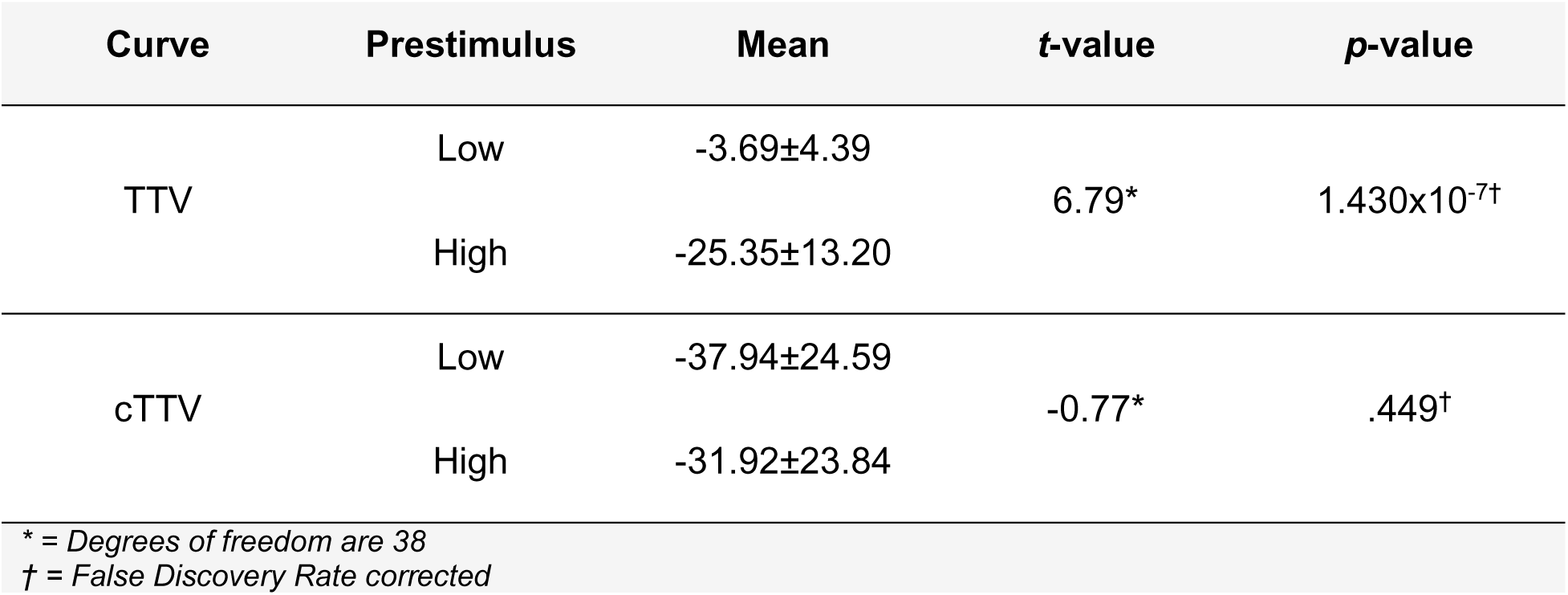
sEEG Maximum quenching in TTV and cTTV repeated measures t-test results

| Curve | Prestimulus | Mean | t-value | p-value |
| --- | --- | --- | --- | --- |
| TTV | Low | -3.69±4.39 | 6.79* | 1.430x10<sup>-7†</sup> |
|  | High | -25.35±13.20 |  |  |
| cTTV | Low | -37.94±24.59 | -0.77* | .449† |
|  | High | -31.92±23.84 |  |  |

\* = Degrees of freedom are 38
† = False Discovery Rate corrected

**Supplementary Table 3:** Maximum quenching between TTV and cTTV in all trials repeated measures t-test results

| Curve for all trials | Mean | t-value | p-value |
| --- | --- | --- | --- |
| TTV | -18.96±11.38 | 3.23* | .004† |
| cTTV | -34.93±18.94 |  |  |

\* = Degrees of freedom are 38
† = False Discovery Rate corrected

**Supplementary Table 4:**
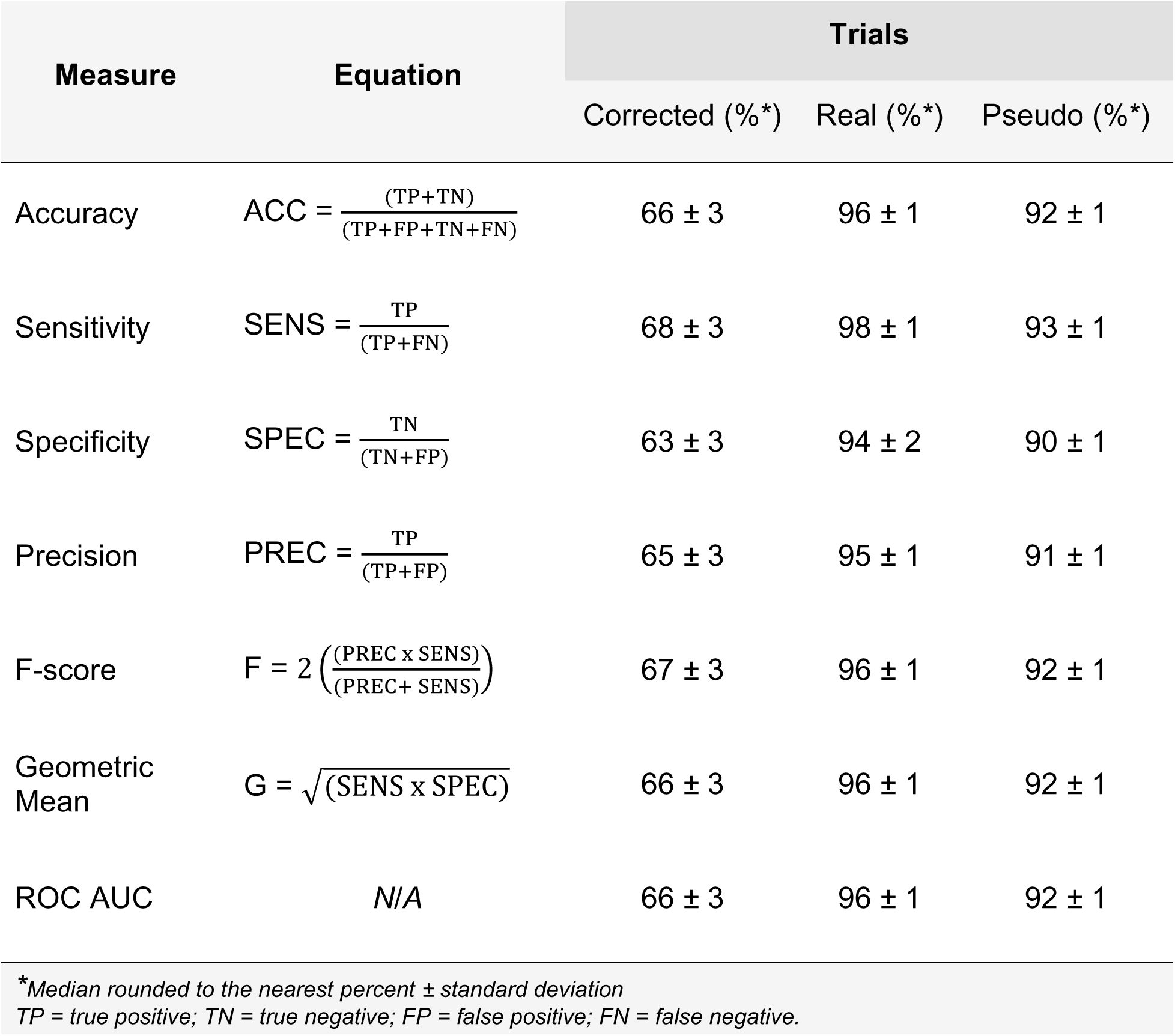
LSTM Deep Learning Classification performance measures

**Supplementary Table 5:** EEG maximum quenching in TTV and cTTV repeated measures t-test results

| Curve | Prestimulus | Mean | *t*-value | *p*-value |
| --- | --- | --- | --- | --- |
| TTV | Low | -9.71±7.69 | 5.59* | 6.193x10<sup>-6†</sup> |
|  | High | -23.91±8.36 |  |  |
| cTTV | Low | -27.43±18.07 | .09* | .930† |
|  | High | -27.87±13.08 |  |  |
| * = Degrees of freedom are 38<br>† = False Discovery Rate corrected |  |  |  |  |

**Supplementary Table 6:** EEG maximum quenching between TTV and cTTV in all trials repeated measures t-test results

| Curve for all trials | Mean | t-value | p-value |
| --- | --- | --- | --- |
| TTV | -16.21±7.64 | 2.31* | .040† |
| cTTV | -23.72±12.40 |  |  |
| * = Degrees of freedom are 38<br>† = False Discovery Rate corrected |  |  |  |

## Notes

### Competing Interest Statement

The authors have declared no competing interest.

